# Endothelial-zippering proceeds by sensing heartbeat-driven force through Cadherin-6 during heart-vessel connection in zebrafish

**DOI:** 10.1101/2025.08.13.669563

**Authors:** Moe Fukumoto, Haruko Watanabe-Takano, Hajime Fukui, Ayano Chiba, Keisuke Sako, Hiroyuki Nakajima, Naoki Mochizuki

## Abstract

The connection between the heart and great vessels established during embryogenesis is essential for circulation. However, how great veins adhere to the endocardium lining the inside lumen of the beating heart remains unknown. Here, using zebrafish, we demonstrate that the endocardium and great veins are sealed in a zipper-closing manner outside the beating heart. The gradual elongation of the endocardium, driven by convergent extension, organized this adhesion by pulling venous ECs along the anterior-posterior axis. Time-specific manipulation of the heart rate revealed that this endocardial elongation proceeds against heartbeat-driven force. From time-lapse imaging of adherens junctions, which would counterbalance mechanical forces, we found a specific contribution of Cadherin-6 instead of Cadherin-5 in sensing endocardium-specific mechanical force. This specificity was confirmed by the depletion of Cadherin-6 that caused endocardium deformation. Altogether, we propose Cadherin-6 mediated EC-zippering as a novel mechanism that updates the understanding of Cadherin usage in dynamic morphogenesis.

## INTRODUCTION

Coordinated development of the heart and blood vessels is essential to establish the closed circulatory system. Previous studies using various model organisms have demonstrated the molecular mechanisms underlying complex heart morphogenesis^1–3^ and vascular network formation.^4,5^ Although the heart and vessels are continuously lined by endothelial cells (ECs), little is known about how vascular ECs adhere to endocardial ECs. Zebrafish are a suitable model to visualize the dynamics of cardiovascular development. The common cardinal vein (CCV) that forms a multicellular sheet-like structure migrates from the dorsal side to the ventral side to connect to the endocardium, which also forms a sheet-like structure outside the beating heart.^6^ Interestingly, the heatbeat starts before CCVs connect to the endocardium. Blood cells gush out of the vein, flow through areas without blood vessels, and get back to the heart.^7^ Although the migrating process of CCV mediated by blood flow has been studied, how CCVs connect to the endocardium remains to be described.

Sealing the gap between two separated tissues is a crucial process during embryogenesis. Zippering is one of the typical sealing systems whereby fusion between opposing epithelia progresses over significant distances. This collective adhesion could be observed during neural tube closure,^8^ optic fissure closure,^9,10^ and tracheoesophageal foregut division.^11^ Diverse morphological and molecular features have been reported in different zippering models, such as filopodia elongation toward opposing cells,^12^ cadherin localization at the adhesive site,^13,14^ and Integrin accumulation with extracellular matrix at the zipper front.^15^ In each model, it is crucial to identify the mechanical force that drives the sequential adhesion to understand the phenomenon in combination with investigating molecular functions.

Cadherin-6, also known as Kidney (K)-Cadherin, is reported to participate in kidney development,^16^ neuron formation,^17^ neural tube closure,^18^ and neural crest cell migration.^19^ As with other typeⅡ Cadherins, the phenotype of Cadherin-6 knockout animals is mild, which makes it difficult to elucidate its function. An important clue to understanding Cadherin’s function is to visualize their localization pattern. For example, Cadherin-5 (Vascular endothelial cadherin; VE-Cadherin), one of the best-studied Cadherin, is reported to form various structures such as linear-^20^, reticular-^21^, junction-associated intermittent lamellipodia (JAIL),^22^ and Cadherin-fingers^23^ that reflect the mechanical condition of the adhesion site.^24,25^ However, visualizing the dynamics of endogenous Cadherins *in vivo* is still a big challenge. Moreover, although Cadherins have been classically identified as tissue-specific molecules,^26^ recent updates in gene expression analysis, such as single-cell RNA-sequencing (scRNA-seq.) revealed mosaic expression patterns of typeⅡ Cadherins across various tissues. ^27^ Thus, considering the combinational and complemental role of different Cadherins in the same tissues or cells has become critical, as well as understanding the function of each Cadherin.

In this study, we demonstrate the contribution of Cadherin-5 and Cadherin-6 in endothelial cell (EC)-zippering during zebrafish heart-vein connection. We first show that CCVs and the endocardium are sealed in a zipper-closing manner. This EC-zippering was facilitated by the constant elongation of the endocardium along the anterior-posterior axis against heartbeat-driven force. Cadherin-6, but not Cadherin-5, is required for endocardial elongation to maintain its sheet structure under complex mechanical conditions.

## RESULTS

### Common cardinal veins adhere to the endocardial sheet at the inflow-tract

In zebrafish, venous blood returns to the heart through the common cardinal veins that show a sheet-like structure (CCV-sheet). ECs of CCV exhibit collective migration from the dorsal toward the ventral to connect to the heart (Figure 1A). The heart starts beating from 24 hours post fertilization (hpf), although the CCVs have not yet connected to the heart.

**Figure 1.**
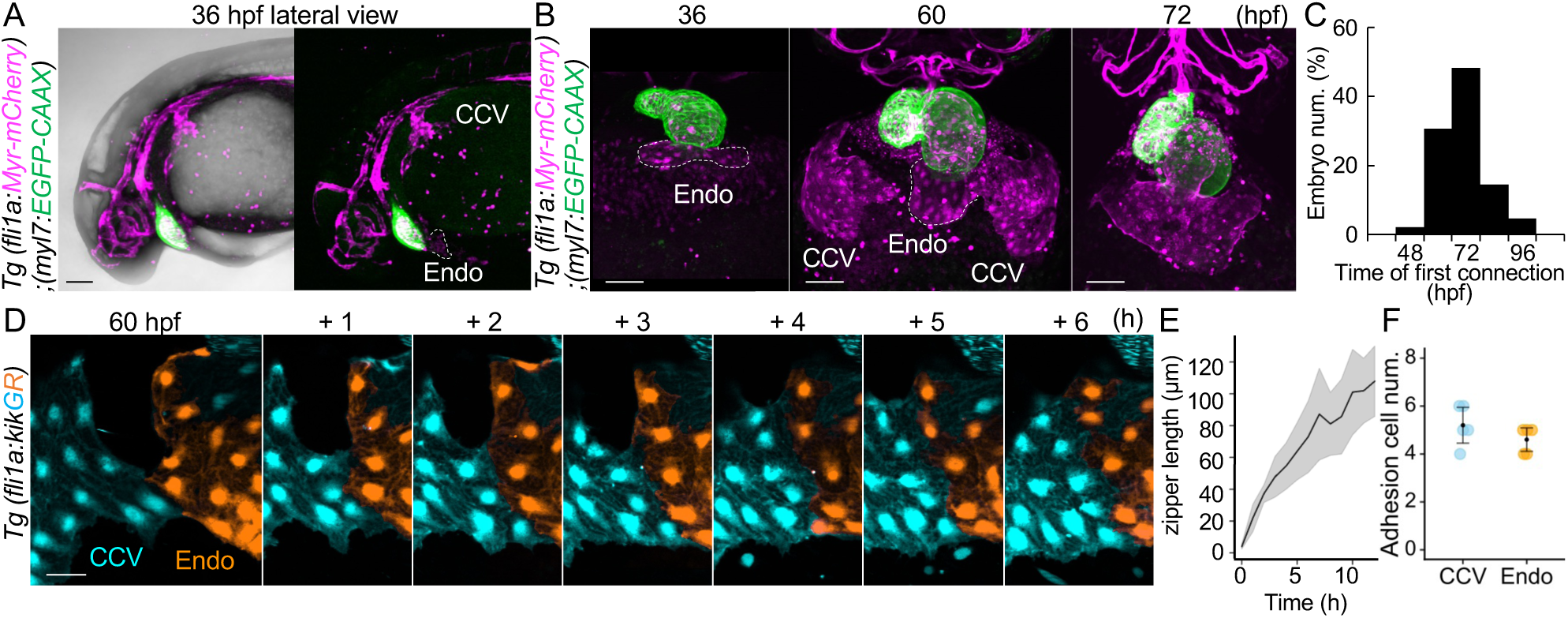
Endo-sheet and CCV-sheet are sealed in a zipper-closing manner. (A) Representative confocal images of *Tg(fli1:Myr-mCherry);(myl7:GFP-CAAX)* embryo. A lateral view of the embryo at 36 h post-fertilization (hpf). The dashed line outlines the endocardium at the inflow-tract (Endo). Images displayed in all the figures are 3D-rendered confocal images of a stack unless otherwise noted. Lateral views, anterior to the right. Ventral views, anterior to the top. Confocal images in all figures are representative of at least three individuals. (B) Representative confocal images of the ventral view of *Tg(fli1:Myr-mCherry);(myl7:GFP-CAAX)* embryo at 36 hpf (left), 60 hpf (center), and 72 hpf (right). The dashed line outlines the endocardium at the inflow-tract (Endo) outside the heart. (C) Quantitative analysis data shown in (B). The histogram shows the percentage of CCV-Endo-connected embryos. *n*=193. (D) Representative time-sequential images of *Tg(fli1a:kikGR);(kdrl:GFP)* embryo from 60 hpf. Photoconverted Endo-sheet (orange) and CCV-sheet (cyan) adhere from one edge to the other in a zipper-closing manner. (E) Quantitative analysis of data shown in (D). The border length of the Endo-sheet and CCV-sheet was measured over time. Data are mean ± s.d.. *n*=5. (F) Quantitative analysis of data shown in (D). Numbers of kdrl:GFP+ ECs on the adhesive front at the end of zippering are shown. Data are mean ± s.d.. *n*=5. Scale bars 100 µm (A, B), 30 µm (D); Endo, endocardium; CCV, common cardinal vein. See also Figures S1 and S2; Videos S1-S7.

To examine the structural connection of the beating heart and CCVs, we observed ECs and cardiomyocytes using *Tg(fli1a:Myr-mCherry);(myl7:GFP-CAAX)*. At the inflow-tract (IFT) of the beating heart, we found an endothelial cell (EC)-sheet posterior to the heart tube at 36 hpf (Figure 1B, left, dashed line). Since this extracardiac EC-sheet could be observed from 22 hpf at the dorsal side continuous to the endocardial ECs (Figure S1A, dashed line), we named this ‘the endocardial sheet (Endo-sheet)’. Bilateral CCV-sheets started attaching to this Endo-sheet around 60 hpf (Figure 1B, center), then CCV-sheets and Endo-sheet were completely fused by about 72 hpf (Figure 1B, right).

To quantify the frequency of heart-CCV connection, we counted the number of embryos that completed the connection. As the histogram showed a wide-range unimodal pattern, there was a large time variation in heart-CCV connection (Figure 1C) compared to the timing of the heart-aorta connection, which occurs at 20 hpf (Figure S1B). Since the majority (49%) of the heart-CCV connection occurred during 60 - 72 hpf, we focused on this time point.

### Endo-sheet and CCV-sheet are sealed in a zipper-closing manner

To understand how the Endo-sheet and CCV-sheets connect, we used *Tg(fli1a:KikumeGR)* line to distinguish each sheet by photo-conversion at 60 hpf (Figure 1D, left). From the time-lapse imaging of the photo-converted embryo, we found that the Endo-sheet and CCV-sheet were sealed in order from one edge to the other (Figure 1D; Video S1). Because similar sequential adhesion modes reported in epithelial tissues are termed epithelial-zippering,^28^ we named this sequential adhesion mode ‘endothelial cell (EC)-zippering’. It took around 10 hours to completely seal the gap between the Endo-sheet and CCV-sheet (Figure 1E). Four to five ECs of each sheet constituted 100 µm in EC-zippering (Figure 1E and 1F). Although there were three patterns in the zippering direction: from posterior to anterior (Video S2), anterior to posterior (Video S3), and both directions toward the center (Figure S2A; Videos S4 and S5), EC-zippering was observed in more than 90% of the embryos from 60 hpf and onwards (Figures S2C and S2D; Video S6). By clear contrast, EC-zippering was not observed in early connected embryos (Figures S2B and S2D; Video S7). These observations indicate that EC-zippering is organized by a developmental stage-related mechanism.

### Endocardial-ECs pull CCV-ECs during EC-zippering

In epithelial zippering, cells move symmetrically to close the gap.^28,29^ To understand how EC-zippering is organized, we examined whether the Endo-sheet and CCV-sheet evenly get close to each other. We first focused on the moving direction and cell shape of the initial adhesion pair. The individual ECs were analyzed using *Tg(fli1a:LIFEACT-mCherry)*, which allowed us to observe the contours of both endocardial-EC (endoEC) and CCV-EC (ccvEC) (Figure 2A; Videovie S8). While ccvEC moved straight towards the endoEC (Figure 2B, left), endoEC temporarily extended its protrusion towards the ccvEC and then returned to its original position (Figure 2B, right). Kymographs revealed that the ccvEC-endoEC border moved approximately 3 µm towards the endoEC side within three hours of initial adhesion (Figure 2C), indicating that EC-zippering is an asymmetric sealing process. Next, we examined whether endoECs pull ccvECs toward the midline or whether ccvECs push endoECs to the midline by losing the EC-sheet tension using single-cell laser-ablation. When we laser-ablated a single endoEC at the posterior side, ccvEC and endoEC recoiled immediately (Figure 2D, left), while ccvEC-endoEC adhesion was maintained when a single ccvEC was ablated (Figure 2D, bottom panel). These results indicate that Endo-sheet tension is required for pulling ccvECs toward the midline at the beginning of EC-zippering.

**Figure 2.**
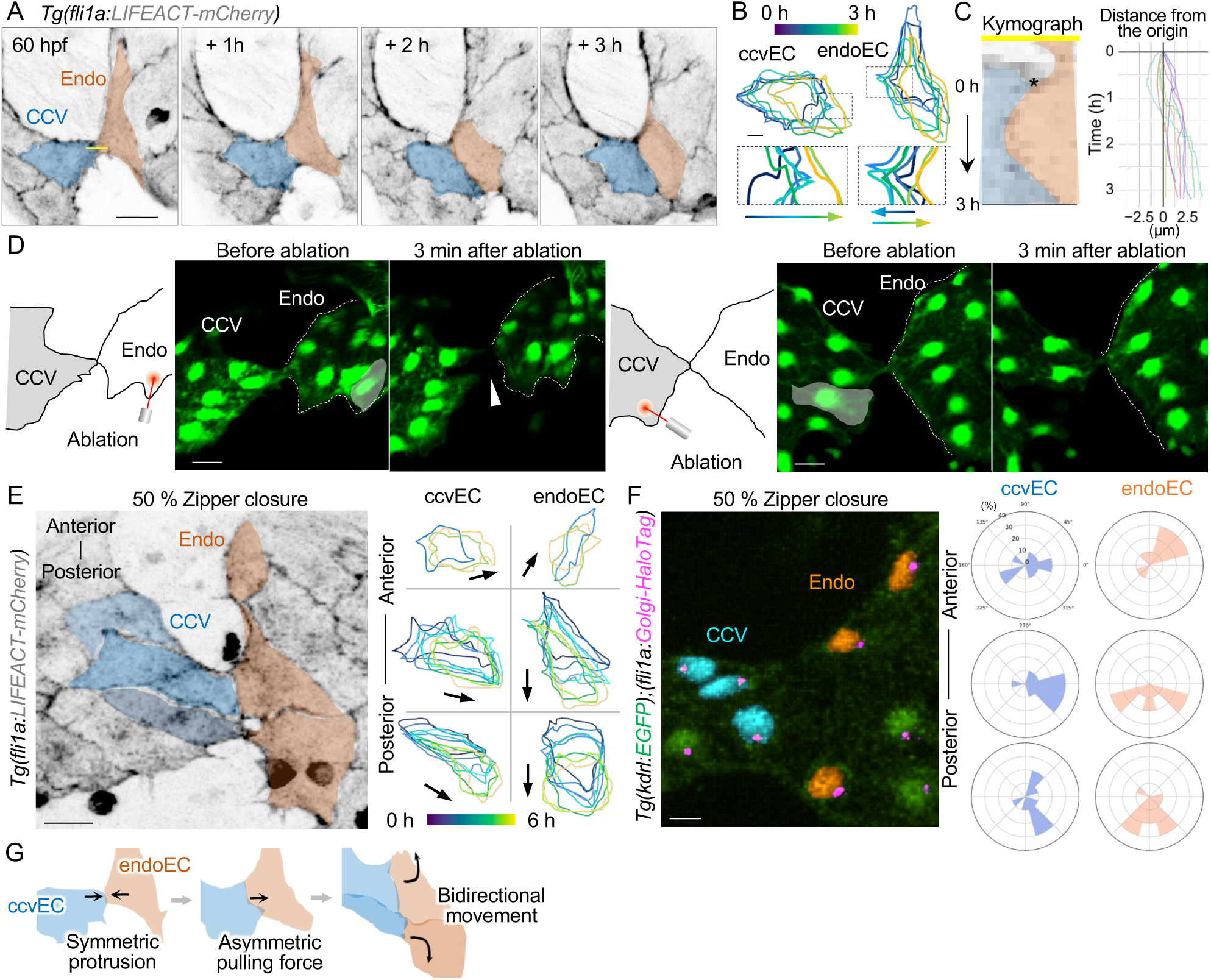
Endocardial-ECs (endoECs) pull CCV-ECs (ccvECs) during EC-zippering. (A) Representative time-sequential images of *Tg(fli1a:LIFEACT-mCherry)* embryo from 60 hpf. The first adhesion pair ECs are labeled in blue (ccvEC) and orange (endoEC). (B) Temporally colour-coded outline at 30-minute intervals of ccvEC (left) and endoEC (right) highlighted in (A). The dashed line box is enlarged (bottom). Color-coded arrows represent the moving direction of the adhesion front. (C) Kymograph at the yellow line from (A). Blue and orange pseudo color represent ccvEC and endoEC, respectively. The position of ccvEC-endoEC boundary was tracked from the initiation of adhesion (labelled by *) for 3 hours and plotted (right). Tracks of 9 embryos were shown in different colors. (D) Schematic images of laser ablation (left). The posterior endoEC (masked white) was laser-ablated at the beginning of EC-zippering (top center). The white arrowhead points to the disconnection of endoEC-ccvEC adhesion after ablation (top right). The posterior ccvEC (masked white) was laser-ablated at the beginning of EC-zippering (bottom center). Dashed lines outline the Endo-sheet. *n*=3 each. (E) Representative image of *Tg(fli1a:LIFEACT-mCherry)* embryo at 63 hpf. Three adhesion pairs were labeled in blue (ccvECs) and orange (endoECs) (left). Temporally color-coded outline at 30-minute intervals of ccvEC and endoEC highlighted in the left image (right). Arrows represent the moving direction of the color-coded outline. (F) Representative image of *Tg(kdrl:EGFP);(fli1a:Golgi-HaloTag)* embryo at 63 hpf. The nuclei of three adhesion pairs were labelled in blue (ccvECs) and orange (endoECs) (left). Quantitative cell population accoring to the cell polarity determined by the cell polarity of Golgi-to-nucleus axis (right). *n*=10. (G) Schematic image of cell movement during EC-zippering. Scale bars, 20 µm (A, D, E, F), 10 µm (B); Endo, endocardium; CCV, common cardinal vein; endoEC, endocardial endothelial cell; ccvEC, common cardinal vein endothelial cell. See also Figure S3; Video S8.

To understand how both EC-sheets sequentially adhere along the posterior-anterior axis, we analyzed the moving direction of the second and third adhesion pairs of ccvEC and endoEC (Figure 2E, left). The posterior endoECs dynamically moved toward the posterior while anterior endoECs slightly moved toward the anterior, whereas ccvECs followed their partner endoECs (Figure 2E, right). Golgi-to-nucleus axis showed a clear bipolar pattern in endoECs and adhered ccvECs (Figure 2F), whereas unadhered ccvEC showed a randomized cell polarity pattern (Figure 2F left-top). To examine whether endoECs pull ccvECs toward the anterior to zip up the zipper, the ablation experiment was conducted in 50% zipped embryos (Figure S3). When a single endoEC at the anterior was ablated, the zipper front was slightly opened, although ccvEC-endoEC adhesion was retained (Figure S3, left panel). There was no difference when anterior ccvEC was ablated (Figure S3, right panel), suggesting that zipper progression toward the anterior also requires Endo-sheet tension that pulls ccvECs toward the anterior. Altogether, these results demonstrate that EC-zippering is an asymmetric zippering, which could be driven by a pulling force generated by the Endo-sheet toward the midline along the anterior-posterior (A-P) axis of the embryo (Figure 2G).

### Endo-sheet changes its shape along the A-P axis

To elucidate how the Endo-sheet produces the pulling force toward the midline together with either anterior or posterior direction, we observed the entire Endo-sheet shape over time. Using *Tg(fli1a:KikumeGR)* line, we photo-converted the endoECs of the Endo-sheet at 2 days post fertilization (dpf) (Figure 3A). The KikumeRed^+^ Endo-sheet showing a horizontal shape spreading along the left-right axis at 2 dpf, gradually narrowed along the left-right axis and extended along the A-P axis, changing into a vertical shape at 3.5 dpf (Figure 3A). Notably, this dynamic transformation was initiated at 2.5 dpf (Figure 3B), the same time-point as EC-zippering was observed, suggesting this Endo-sheet transformation could be the time-related factor to generate the pulling force for EC-zippering. To further understand the mechanical force that would drive this endo-sheet transformation, we assessed the dynamics of actomyosin in endoECs during 2-3 dpf. While individual endoECs dynamically changed their shape from a rounded to a spindle-like shape (Figure 3C) with less size (Figure 3D), there were no significant changes in myosin accumulation (Figure 3E). These results suggest that endo-sheet transforms passively, instead of actively changing their shape.

**Figure 3.**
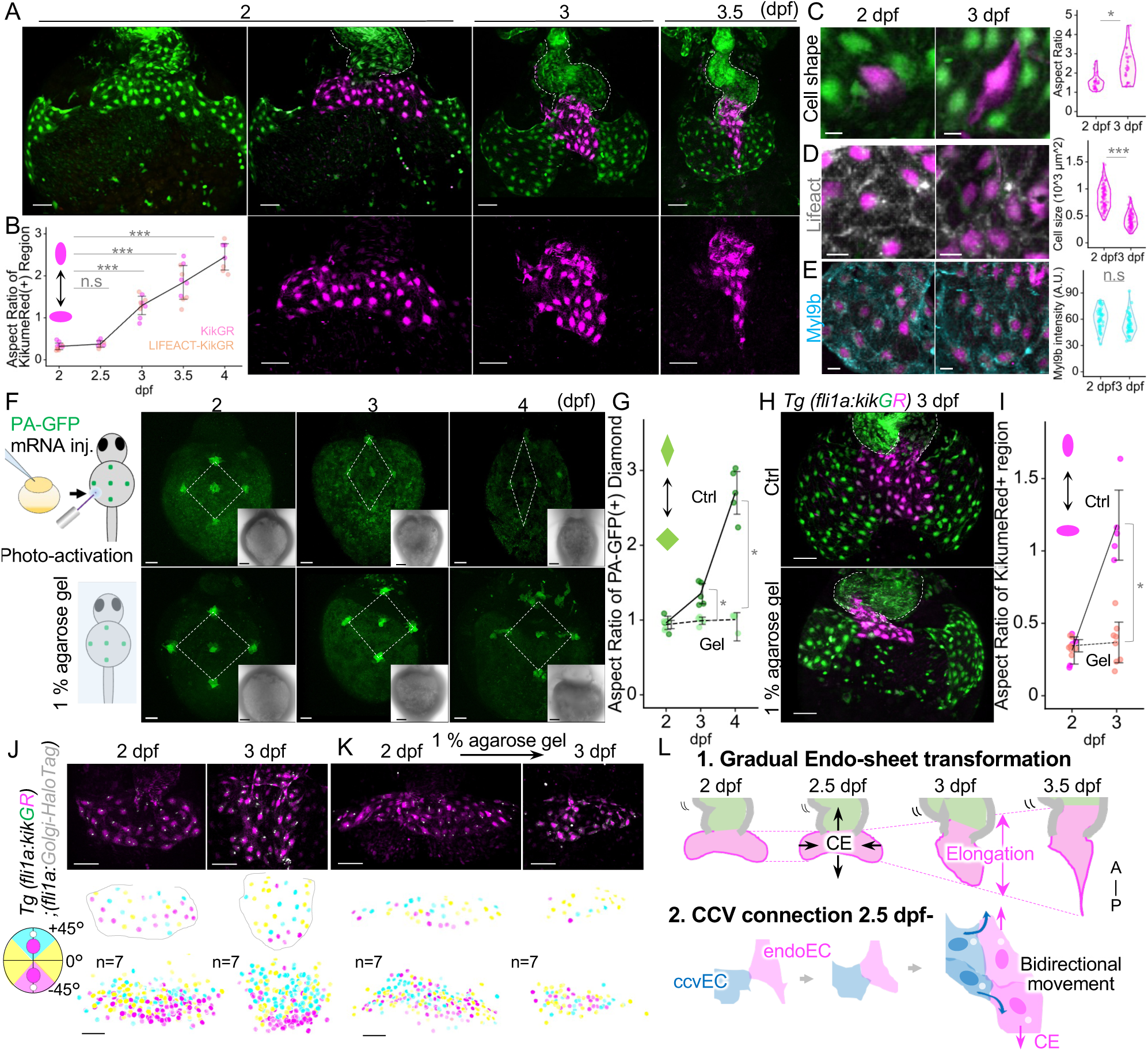
Endo-sheet gradually elongates along the anterior-posterior axis by convergent extension. (A) Representative snap-shot images of *Tg(fli1a:KikumeGR)* embryo from 2 dpf to 4 dpf (top panel). The dashed line outlines the heart. The Endo-sheet photoconverted into magenta is enlarged (bottom panel). (B) Quantitative analysis of the aspect ratio (the Left-right length to the anterior-posterior length ratio) of KikumeRed-positive region was measured. Data are mean ± s.d. *n*=11. (C) High magnification snap-shot images of *Tg(fli1a:KikumeGR)* embryo at 2 dpf and 3 dpf. A single endoEC was photoconverted (magneta, left panel). The aspect ratio (the long-to-short length ratio) was measured and plotted (right). *n*= 32. (D) High magnification snap-shot images of *Tg(fli1a:KikumeGR);(fli1a:LIFEACT-HaloTag)* embryo at 2 dpf and 3 dpf. endoECs were photoconverted at 2 dpf (magneta, left panel). The cell size was measured and plotted (right). *n*= 108. (E) High magnification snap-shot images of *Tg(fli1a:KikumeGR);(fli1a:myl9b-HaloTag)* embryo at 2 dpf and 3 dpf. endoECs were photoconverted at 2 dpf (magneta, left panel). Intensity of HaloTag-positive Myl9b of *Tg(fli1a:myl9b-HaloTag)*. *n*=80. (F) Experimental design of ventral tissue labeling (top left). Representative snap-shot images of five PA-GFP spots on the ventral tissue from 2 dpf to 4 dpf (top panel). Representative snap-shot images of five PA-GFP spots on the ventral tissue from 2 dpf to 4 dpf of 1% agarose gel embedded embryos (bottom panel). Inset shows the bright field image of ventral tissue. (G) Quantitative analysis of data shown in (C). The aspect ratio of PA-GFP labeled diamond was measured from 2 dpf to 4 dpf. Data are mean ± s.d. *n*=5. (H) Representative images of *Tg(fli1a:KikumeGR)* embryo of gel-free (top) and gel-embedded (bottom) embryo at 3 dpf. The dashed line outlines the heart. (I) Quantitative analysis of data shown in (E). The aspect ratio of KikumeRed positive region was measured. Data are mean ± s.d. Ctrl, *n*=5; Gel, *n*=8. (J) Representative snap-shot images of *Tg(fli1a:KikumeGR);(fli1a:Golgi-HaloTag)* embryo at 2 dpf and 3 dpf. endoECs were photoconverted into magenta at 2 dpf (top). The Golgi-to-nucleus axes of each endoEC were measured and mapped as a pseudocolour representation of the nucleus (middle). Additive projections of the cells exhibiting the polarity difference anterior-(cyan), posterior-(magenta), and other direction (yellow) at two different time point (2 dpf and 3 dpf). *n*=7 (bottom). (K) Representative snap-shot images of *Tg(fli1a:KikumeGR);(fli1a:Golgi-HaloTag)* embryo embeded in 1% agarose gel at 2 dpf and 3 dpf. endoECs were photoconverted at 2 dpf (magneta, top). Additive projections same as (J) *n*=6 (bottom). (L) Schematic image of Endo-sheet transformation. Endo-sheet gradually elongates from 2.5 dpf and gains bipolarity at 3 dpf. Scale bars, 10 µm (C, D, E), 50 µm (A, F, H, J, K) and 100 µm (inset C); PA, photoactivatable. Statistical analysis was performed by the paired samples *t*-test (A, C) and Welch’s *t*-test (G, I).*p<0.05, ***p<0.01. See also Figures S4 and S5.

### Embryonic elongation pulls the Endo-sheet toward the posterior

During embryogenesis, embryos transform spontaneously along the body axis, which is organized by a collective behavior called convergent extension (CE). ^30,31^ From 2 dpf to 4 dpf, ventral tissue around the heart gradually changed its shape following the typical CE mode (Figure S4A). To investigate whether Endo-sheet transformation is caused by CE, we examined the movement of ventral cells labeled by photoactivation. Photo-activatable (PA)-GFP-labelled cells were observed over time (Figure 3F, illustration). The ventral tissue gradually followed a typical CE mode from 2 dpf to 4 dpf as PA-GFP+ cells along the left-right axis moved toward the midline, and those along the A-P axis moved away along the A-P axis (Figure 3F, top panels, and 3G). Since we knew that agarose-gel embedding for long-time-lapse imaging impairs embryonic elongation and causes ventral tissue deformation (Figure S4B), we examined if agarose-gel embedding inhibits CE. As expected, the CE mode was completely inhibited when the embryos were embedded in 1% agarose gel from 2 dpf to 4 dpf (Figures 3F, bottom panels, and 3G). In agarose gel-embedded embryos, KikumeRed^+^ Endo-sheet failed to elongate toward the posterior side (Figures 3H and 3I), indicating that CE drives Endo-sheet transformation. To elucidate the relationship between endo-sheet transformation and the bidirectional movement of endoECs demonstrated in Figures 2E and 2F, we analyzed EndoEC polarity during 2-3 dpf by the position of Golgi in cells. At 2 dpf, cell polarity was randomized. By contrast, at 3 dpf, cells at the posterior edge faced toward the posterior side, whereas cells at the anterior side faced toward the anterior side. (Figure 3J). When CE was inhibited by gel embedding, Endo-sheets failed to gain this position-dependent polarity (Figure 3K). Collectively, these results indicate that CE induces the Endo-sheet elongation accompanied by the EC position-dependent bipolarity.

### Endo-sheet elongation is inhibited by the heartbeat

KikumeRed^+^ endoECs were detected inside the atrium from 3 dpf (Figure 3A), even in 1% agarose gel-embedded embryos (Figure 3H). Therefore, we hypothesized that another force related to heart dynamics may affect Endo-sheet transformation toward the anterior. We examined whether heartbeat (HB) affects the Endo-sheet transformation by treating embryos with verapamil, which reduces HB. In verapamil-treated embryos, the Endo-sheet elongated toward the posterior. Conversely, when HB was increased by isoprenaline treatment, the Endo-sheet shrank and shifted toward the anterior. The elongation caused by verapamil treatment was canceled when CE was inhibited (Figures 4A and 4B). These results indicate that the Endo-sheet is elongated by CE against HB-driven force.

**Figure 4.**
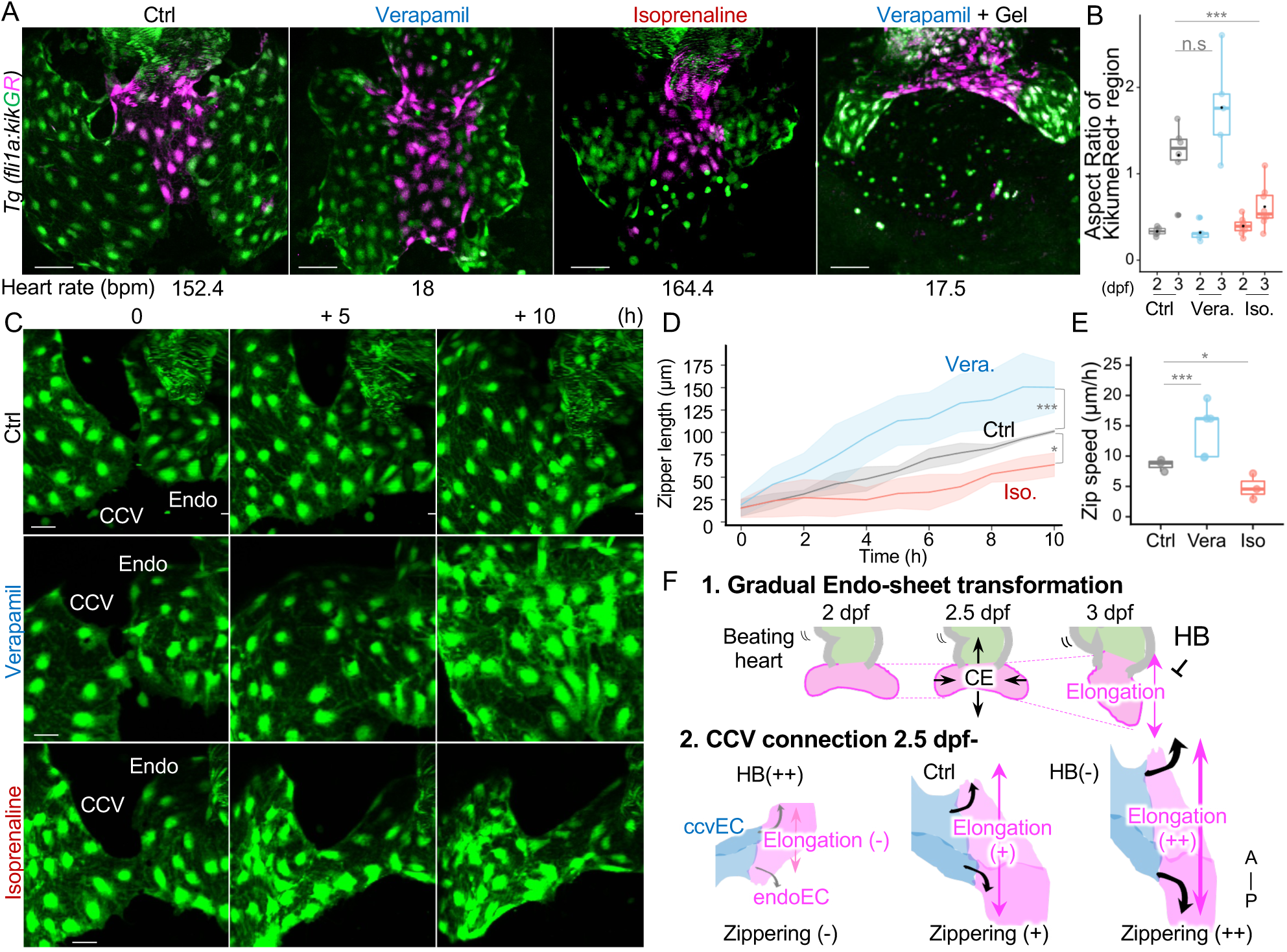
Heartbeat inhibits End-sheet elongation and EC-zippering. (A) Representative images of *Tg(fli1a:KikumeGR)* embryo at 3 dpf treated with DMSO, 200 µM verapamil, 300 µM isoprenaline hydrochloride, and 200 µM verapamil in gel embedded condition for 24 h. Heart rates are mean. *n*=3. (B) Quantitative analysis of data shown in (B). The aspect ratio of KikumeRed positive region was measured. Data are mean ± s.d. *n*=5-8. (C) Representative time-sequential images of *Tg(kdrl:EGFP)* embryos from 60 hpf. Embryos were treated with DMSO (top), 200 µM verapamil (middle), and 300 µM isoprenaline hydrochloride (bottom). (D) Quantitative analysis of data shown in (C). Length of kdrl:GFP positive region from the first adhesion point to the zipper front was measured. Data are mean ± s.d. *n*=3. (E) Average zippering speed shown in (D). Data are mean ± s.d. *n*=3. (F) Schematic image of EC-zippering mechanism. 1. Endo-sheet transformation occurs from 2.5 dpf. 2. As the Endo-sheet elongates against HB-driven force, ccvECs are pulled bidirectionally along the A-P axis. Scale bars, 50 µm (A), 20 µm (C); Iso, isoprenaline; Vera, verapamil; HB, heartbeat; CE, convergent extension; endoEC, endocardial endothelial cell; ccvEC, common cardinal vein endothelial cell; n.s, no significance. Statistical analysis was performed by the Welch’s *t*-test (B, D, E). *p<0.05, ***<0.01. See also Videos S9-S11.

### EC-zippering proceeds against heartbeat

To clarify whether Endo-sheet transformation generates the pulling force for EC-zippering, we promoted Endo-sheet elongation by treating embryos with verapamil from the beginning of EC-zippering (Figure 4C, middle panels; Video S9). Zippering length and speed were increased and accelerated, respectively, in verapamil-treated embryos (Figures 4D and 4E; Videos S10 and S11). By contrast, when we induced Endo-sheet shrinkage toward the anterior by isoprenaline treatment, EC-zippering was completely impaired (Figure 4C, bottom panels, 4D, and 4E), demonstrating that EC-zippering can be controlled only by manipulating the heart rate that imbalances the Endo-sheet morphology. These results indicate that EC-zippering is facilitated by Endo-sheet elongation that proceeds against HB (Figure 4F).

### Cadherin-6 is an EC-sheet-specific adhesion molecule

Since Endo-sheets appeared to receive complex mechanical forces produced by CE and HB, we assumed the presence of Endo-sheet-specific intercellular adhesion molecules that balance the forces between HB and CE to maintain the Endo-sheet structure during development. We searched for such intercellular adhesion molecules by re-analyzing a single-cell RNA sequence data set^32^ and identified Cadherin-6 as a candidate (Figure S5).

To examine whether Cadherin-6 is exactly expressed in EC-sheets, we established Cdh6-EGFP knock-in zebrafish line, *TgKI(cdh6-EGFP)*, in which coding sequence of enhanced GFP (EGFP) was inserted at the C terminus of endogenous Cadherin-6, using CRISPR-Cas9 system (Figure 5A). Cdh6-EGFP signal was observed along the cell-cell junctions of the spinal cord, floorplate, hypochord, and veins, consistent with the scRNA-seq data (Figure S6). At the IFT, Cdh6-EGFP was detected in the CCV-sheets and Endo-sheet but not in the endocardium lining the inside lumen of the heart (Figure 5B), suggesting its significance in EC-sheets.

**Figure 5.**
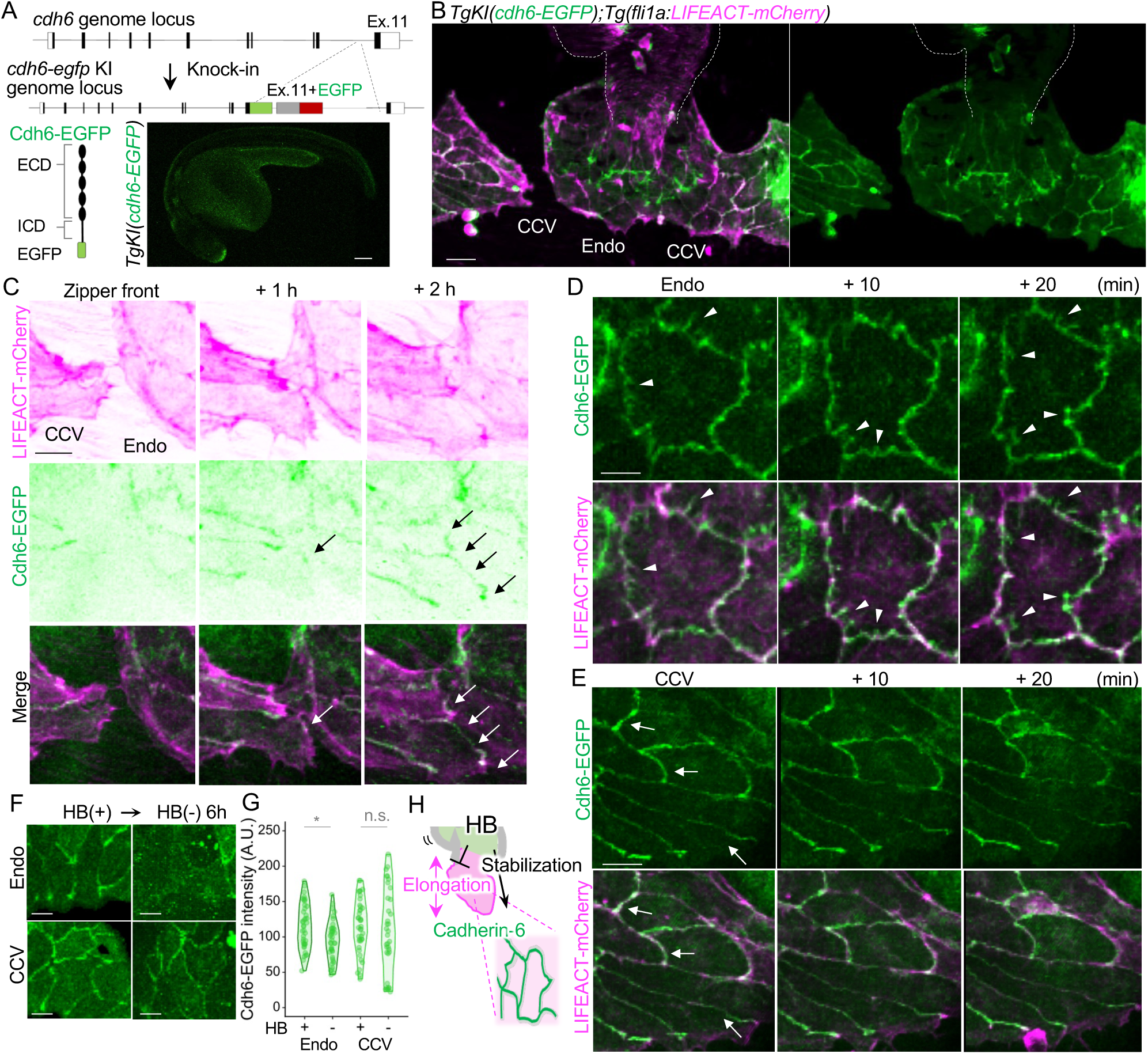
Cadherin-6 localization in EC-sheets. (A) Schematic image of *TgKI(cdh6-EGFP)* development. EGFP was inserted after the last exon of *cdh6* to visualize endogenous Cdh6-EGFP fusion protein (top). Representative confocal images of the lateral view of *TgKI(cdh6-EGFP)* embryo at 24 hpf (bottom). (B) Representative confocal images of the *TgKI(cdh6-EGFP);Tg(fli1a:LIFEACT-mCherry)* line at 60 hpf. The dashed line outlines the heart. (C) High magnification time sequential confocal images of the *TgKI(cdh6-EGFP);Tg(fli1a:LIFEACT-mCherry)* embryo at the adhesion front of CCV-sheet and Endo-sheet from 63 hpf. Arrows point to the emerging Cdh6-EGFP signal at the adhesion front. (D) High magnification time sequential confocal images of the *TgKI(cdh6EGFP);Tg(fli1a:LIFEACT-mCherry)* embryo at the Endo-sheet from 60 hpf. Arrowheads point to the protrusion-like Cdh6-EGFP. (E) High magnification confocal images of the *TgKI(cdh6-EGFP);Tg(fli1a:LIFEACT-mCherry)* embryo at CCV at 60 hpf. Arrows point to the Cdh6-EGFP linearized signals at the cell-cell adhesion site. (F) High magnification confocal images of the *TgKI(cdh6-EGFP);Tg(fli1a:LIFEACT-mCherry)* embryo at Endo-sheet (top) and CCV-sheet (bottom) before and 6 hours after Verapamil treatment around 60 hpf. (G) Quantitative analysis of data shown in (F). The Cdh6-EGFP signal intensity was measured. *n*=40. (H) Schematic image of Cadherin-6 expression on Endo-sheet. Scale bars, 100 µm (A), 50 µm (B),10µm (C),15 µm (D and E); Endo, endocardium; CCV, common cardinal vein. See also Figure S6-S8 ; Videos S12-S14.

### Cadherin-6 localization dynamically changes during EC zippering

In addition to the expression pattern in tissues, subcellular localization reflects the function of Cadherins. To understand the role of Cadherin-6 in EC-sheets, we closely looked at the localization of Cdh6-EGFP with actin cytoskeleton using *TgKI(cdh6-EGFP);(fli1a:LIFEACT-mCherry)*. At the zipper front, where ccvECs adhered to endoECs, Cdh6-EGFP was slightly detected at the beginning, then EGFP intensity became stronger as the zippering proceeded and was also detected as a linearized signal as adhesion matured (Figure 5C, arrows; Video S12). In Endo-sheets, Cdh6-EGFP was detected as a protrusion-like structure accompanied by actin filaments that resembled the structure called Cadherin fingers (Figure 5D).^23^ We noticed that Cadherin fingers constantly changed direction by taking short-interval time-lapse imaging (Figure 5D, arrowheads; Video S13). By clear contrast, in the CCV-sheets, Cdh6-EGFP was detected as a linearized signal along the cell-cell junctions (Figure 5E, arrows; Video S14). Since we assumed Cadherin-6 as an adhesion molecule that balances the forces between HB and CE in Endo-sheet structure during development, we checked whether HB alters the expression pattern of Cadherin-6. When HB was stopped by verapamil treatment, Cdh6-EGFP signal significantly decreased in endo-sheets, while Cdh6-EGFP signal intensity remained unchanged in CCV-sheet (Figures 6F and 6G). These data support the idea that Cadherin-6 specifically responds to HB-dependent mechanical forces in Endo-sheet (Figure 6H).

**Figure 6.**
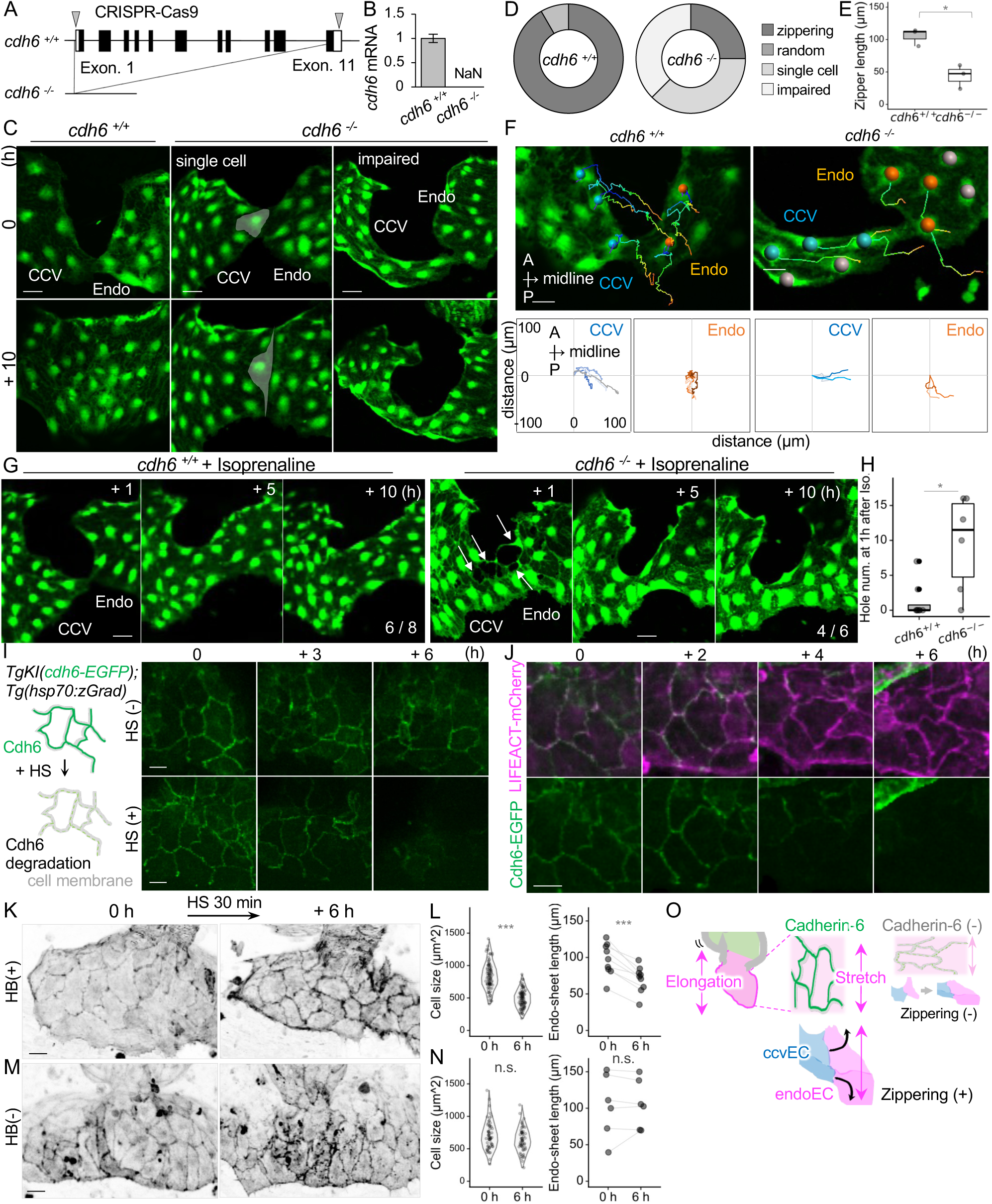
Cadherin-6 is required for EC-zippering through maintaining endoEC size against heatbeat. (A) Schematic image of *cdh6* KO fish genome. The entire *cdh6* genome locus was deleted by CRISPR-Cas9 system. (B) Quantification of *cdh6* mRNA level of 55 hpf embryos. *n*=3. (C) Representative confocal images of the ventral view of *Tg(kdrl:EGFP) cdh6^+/+^* or *cdh6^-/-^* embryo at 60 hpf (top) and 70 hpf (bottom). (D) Quantification of the phenotype shown in (C). *cdh6^+/+^*, *n*=19; *cdh6^-/-^*, *n*=9. (E) Quantitative analysis of data shown in (C). Length of kdrl:GFP positive region from the first adhesion point to the zipper front was measured at 70 hpf in *cdh6^+/+^* embryos or *cdh6^-/-^* embryos showing zipper defect. Data are mean ± s.d. *n*=3. (F) Confocal images analysing the moving direction of EC adhesion pairs in *cdh6^+/+^* and *cdh6^-/-^* embryos shown in (C). endoECs (orange) and ccvECs (blue) were labelled and tracked over time (top). Quantitative analysis of displacement length and direction of endoECs and ccvECs (bottom). (G) Representative time-sequential image of *Tg(kdrl:EGFP) cdh6^+/+^* or *cdh6^-/-^* embryos treated with isoprenaline. Arrows point to the weakened cell-cell adhesion region. The frequency of the phenotype is shown at the bottom right. (H) Quantitative analysis of data shown in (G). Number of kdrl:GFP negative region in the Endo-sheet of *cdh6^+/+^* embryos or *cdh6^-/-^* embryos was measured at 1 hour after 300 µM isoprenaline treatment. *cdh6^+/+^*, *n*=8; *cdh6^-/-^*, *n*=8. (I) Schematic image of Cdh6-EGFP degron system (left). Representative time-sequential confocal images of *TgKI(cdh6-EGFP);Tg(hsp70:zGrad)* embryos without heat shock (top) and with heat shock (bottom). (J) Time-sequential images of *TgKI(cdh6-EGFP);Tg(hsp70:zGrad);(fli1a:LIFEACT-mCherry)* embryo after heat shock. (K) Representative Endo-sheet images of *TgKI(cdh6-EGFP);Tg(hsp70:zGrad);(fli1a:LIFEACT-mCherry)* line soon after heat shock (left) and 6 h after heat shock (right). (L) Quantitative analysis of data shown in (I). Individual cell sizes outlined by LIFEACT-mCherry are shown. *n*=90. Endo-sheet lengths along the A-P axis before and after heat shock are shown. *n*=9. (M) Representative Endo-sheet images of *TgKI(cdh6-EGFP);Tg(hsp70:zGrad);(fli1a:LIFEACT-mCherry)* soon after heat shock (left) and 6 h after heat shock (right) without heatbeat. (N) Quantitative analysis of data shown in (K). Individual cell sizes outlined by LIFEACT-mCherry are shown. *n*=60.Endo-sheet lengths along the A-P axis before and after heat shock are shown. *n*=6. (O) Schematic image of Cadherin-6 mediated EC-zippering. Scale bars, 20 µm (C, F, G, K, M), 10 µm (I and J); NaN, not a number; Endo, endocardium; CCV, common cardinal vein; HS, heat shock; Iso, isoprenaline; Vera, verapamil; HB, heatbeat; CE, convergent extension. Statistical analysis was performed by the Student’s *t*-test (E, H, L-left and N-left) or paired samples *t*-test (L-right and N-right). *p<0.05,***p<0.01. See also Videos S15-S19.

### Cadherin-6 is required for EC-zippering

To clarify the role of Cadherin-6 in Endo-sheets, we established a *cdh6* knock-out (*cdh6^-/-^*) zebrafish line. We deleted the entire *cdh6* genome locus by CRISPR-Cas9 system (Figure 6A) to avoid genetic compensation.^33^ This *cdh6^-/-^*fish completely lacked *cdh6* mRNA (Figure 6B). In *cdh6*^-/-^ embryos, both Endo-sheet and CCV-sheet were successfully formed without any intercellular gaps (Figure 6C). However, EC-zippering was impaired in 60% of the *cdh6*^-/-^ embryos (Figures 6C and 6D). In 30% of embryos, a single elongated EC formed the adhesion between the two sheets (Figure 6C, middle; Videos S15 and S16). Another 30% of embryos showed impaired closure between two EC sheets (Figures 6C, right, and 6E; Video S17). To clarify the reason for zipper defects by deleting *cdh6*^-/-^, we analyzed the moving direction of ccvECs and endoECs. In the control embryos, endoECs moved toward the posterior direction (Figure 6F, top). By contrast, in *cdh6*^-/-^ embryos, endoECs moved toward the midline (Figure 6F, bottom). ccvECs in both control and *cdh6*^-/-^ embryos moved toward their adhesion partner. These results indicate that Cadherin-6 is required for balancing the force in the Endo-sheet. We further confirmed if Cadherin-6 functions under HB-driven mechanical force, which would be specific in the Endo-sheet. When *cdh6*^-/-^ embryos were treated with isoprenaline, EGFP-negative holes were observed at 1 hour after treatment, suggesting that Endo-sheet cell-cell adhesions were transiently weakened in the absence of *cdh6* (Figures 6G and 6H; Videos S18 and S19). These results further support the idea that Cadherin-6 is required to maintain cell adhesions under HB-driven mechanical force.

### Cdherin-6 is indispensable for maintaining the Endo-sheet structure

As *cdh6* deletion affected EC-zippering only in a portion of embryos (Figure 6D) and as isoprenaline treatment to *cdh6^-/-^* embryos only induced a transient defect (Figure 6G), we hypothesized that unspecified factors compensate for the constant lack of Cadherin-6. Therefore, we further directly examined the contribution of Caherin-6 in Endo-sheet force balancing by depleting Cadherin-6 protein at a specific time point. We used a zGrad-based degron system^29^ to degrade Cdh6-EGFP in a heat-shock (HS) inducible way (Figure 6I, illustration). ^34^ In *TgKI(cdh6-EGFP);Tg(hsp70:zGrad)* embryos, the fluorescence of Cdh6-EGFP decreased from 3 hours post-HS and completely disappeared at 6 hours post-HS (Figure 6I, right panels). As Cdh6-EGFP degraded, each endoEC shrank (Figures 6J and 6K), resulting in the shrinkage of the entire Endo-sheet (Figures 6K and 6L). To examine if Cadherin-6 is required to maintain the Endo-sheet structure under HB-driven force, we further temporally degraded Cadherin-6 using the degron system under verapamil treatment. When HB was stopped, individual cell size and the entire Endo-sheet structure were maintained, even Cadherin-6 was degraded (Figures 6M and 6N). These results indicate that Cadherin-6 is indispensable to maintain the Endo-sheet structure by preserving individual cell shape under HB-driven force (Figure 6O).

### Cadherin-5 is dispensable for maintaining the Endo-sheet structure

Cadherin-5 is the most well-studied Cadherin in ECs. To map the significance of Cadherin-6 in EC-sheets, we examined the contribution of Cadherin-5 in Endo-sheet maintenance. To investigate Cadherin-5 expression and localization pattern, we established *TgKI(cdh5-EGFP)* line similar to *TgKI(cdh6-EGFP)*. Cdh5-EGFP was detected along the cell-cell junction of the entire endothelium (Figure 7A), indicating that degradation of Cdh6-EGFP was not rescued by the resident Cadherin-5. Unlike Cadherin-6, Cdh5-EGFP was detected only as a linearized signal with a variable signal intensity in the Endo-sheet (Figure 7B; Video S20). To directly compare the localization pattern of Cadherin-6 and Cadherin-5, we also established *TgKI(cdh6-mCherry)* and crossed it with *TgKI(cdh5-EGFP)*. Although Cadherin-5 and Cadherin-6 colocalized in most cell-cell junctions (Figure 7C, white arrow), complemental localization was observed in specific regions. Cdh5-EGFP could be detected at the widened cell-cell adhesion site (Figure 7C, yellow arrowheads), whereas Cdh6-mCherry preferentially localized at the sharpened cell-cell adhesion site (Figure 7C, white arrowheads). This observation was consistent with the idea that Cadherin-6 respond to mechanical force more sensitively than Cadherin-5, suggesting the functional difference between these two Cadherins.

**Figure 7.**
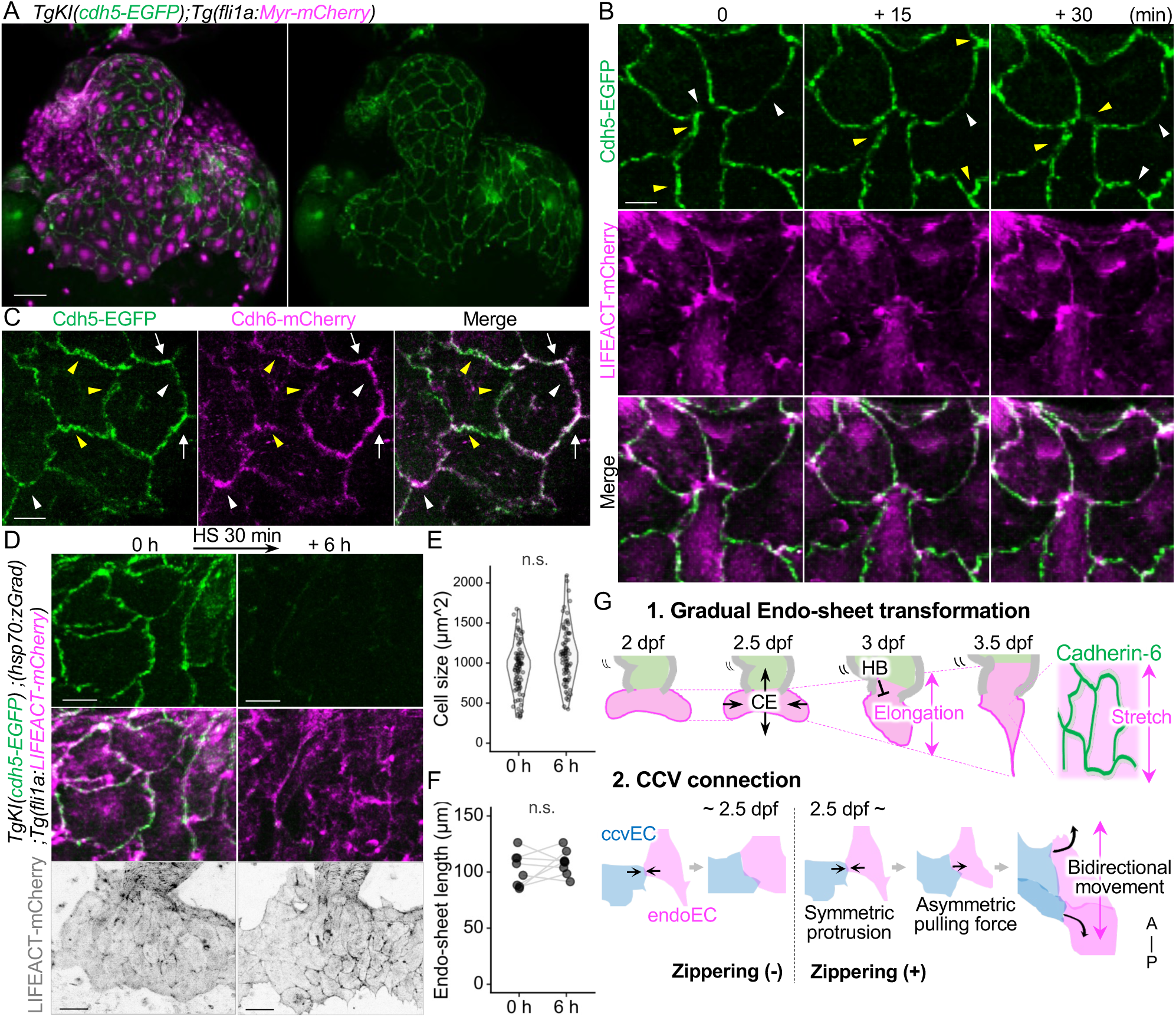
Cadherin-5 is dispensable for maintaining endoEC size and Endo-sheet structure. (A) Representative confocal images of the *TgKI(cdh5-EGFP);Tg(fli1a:Myr-mCherry)* embryo. Ventral view at 60 hpf. (B) High-magnification time-sequential confocal images of Endo-sheet of the *TgKI(cdh5-EGFP);Tg(fli1a:LIFEACT-mCherry)* embryo at 60 hpf. White arrowheads point to Cdh5-EGFP weak or negative cell-cell junctions. Double arrowheads point strong and/or broad Cdh5-EGFP signals. (C) High-magnification confocal images of Endo-sheet of the *TgKI(cdh6-mCherry);(cdh5-EGFP)* embryo. White arrows point to the Cdh5-EGFP and Cdh6-mCherry colocalized sites. White arrowheads point Cdh6-EGFP dominant site. Double arrowheads point Cdh5 dominant site. (D) Representative Endo-sheet images of *TgKI(cdh6-EGFP);Tg(hsp70:zGrad);(fli1a:LIFEACT-mCherry)* line soon after heat shock (top) and 6 h after heat shock (bottom). The entire shape of the Endo-sheets were visualized by LIFEACT-mCherry (right). (E) Quantitative analysis of data shown in (D). Individual cell sizes outlined by LIFEACT-mCherry are shown. *n*=80. (F) Quantitative analysis of data shown in (D). Endo-sheet lengths along the A-P axis before and after heat shock are shown. *n*=8. (G) Schematic image of Cadherin-6 mediated EC-zippering. Endocardial sheet transformation forced by convergent extension and heatbeat pulls the ccvECs along A-P axis. Scale bars, 50 µm (A),10 µm (B, C, D-left), 40 µm (D-right); CCV, common cardinal vein; HB, heatbeat; CE, convergent extension; endoEC, endocardial endothelial cell; ccvEC, common cardinal vein endothelial cell; A, anterior; P, posterior; A-P, anterior-posterior; n.s., no significance. Statistical analysis was performed by the Student’s *t*-test (E) or the paired samples *t*-test (F). See also Video S20.

To examine if Cadherin-5 alone is required for Endo-sheet maintenance, we also used the HS-induced degron system to degrade Cdh5-EGFP using *TgKI(cdh5-EGFP);Tg(hsp70:zGrad)(fli1a:LIFEACT-mCherry)* line. Even though Cdh5-EGFP was degraded at the cell-cell adhesions, endoEC size and Endo-sheet structure were unchanged (Figures 7D, 7E, and 7F). These results emphasize the specific contribution of Cadherin-6 in maintaining cell size and sheet structure of Endo-sheet, which is essential for EC-zippering (Figure 7G).

## DISCUSSION

In this study, we demonstrate the molecular mechanism of IFT formation in zebrafish. We first show that CCV-sheet and Endo-sheet attach by EC-zippering. This collective adhesion required the Endo-sheet elongation along the A-P axis, which was counterbalanced presumably by the forces driven by CE and HB (Figure 7G). At a molecular level, Cadherin-6 expressed in Endo-sheets appeared to sense these mechanical forces sharply. When Cadherin-6 was depleted, individual endoEC shrank. Therefore, the Endo-sheet failed to elongate along the A-P axis, which led to disturbed EC-zippering.

We captured a complemental localization of Cadherin-5 and Cadherin-6 in transforming Endo-sheets. Since Cadherin localizations are affected by mechanical forces they receive,^35^ two structural features in Cadherin-6 considerably explain this difference. One is the hydrophobic amino acid (AA) sequence in the β-catenin binding domain, which might strengthen the bond between Cadherin-6 and β-catenin. The other is the RGD sequence in the first Cadherin domain, which has the potential to bind with Integrins.^36^ Since the role of RGD-domains in Cadherins has not been clarified in development and taken that Cadherin-6 is the only RGD-Cadherin in zebrafish, Endo-sheet could be a suitable model to understand the significance of RGD-Cadherins. In addition to these Cadherin-6 specific AAs, previous studies have reported that Cadherin-6 has the highest homophilic binding affinity among other type-2 Cadherins in mice^37^, which would explain the reason for Cadherin-6 use instead of other Cadherins in dynamic EC-sheets.

Adherens junctions function as crucial sensors to adapt cells to various mechanical conditions.^35,38^ The Cadherin-finger-like structure of Cadherin-6, particularly found in Endo-sheets, implies the specific function of Cadherin-6 in response to strong mechanical forces.^23^ Time-specific Cadherin-6 depletion demonstrated its function in maintaining the cell size. Even though the precise characteristics of the forces driven by heartbeat (e.g., strength and frequency) need to be determined, our study provides a molecular explanation of how cardiovascular tissues maintain their structure under the mechanical influence of continuous heartbeat.

The gradual transformation of the Endo-sheet balanced by Cadherin-6 appeared to facilitate EC-zippering by pulling ccvECs bidirectionally along the A-P axis. This transformation would be the time-related factor for EC-zippering that explains why EC-zippering was undetectable at earlier stages. This morphological diversity presumably imply the complexity and difficulty of the mechanism underlying how CCV connects to the heart. In addition, we found that agarose gel embedding, a general condition for zebrafish time-lapse imaging, would be an artifact for observing EC-zippering by affecting the morphology of EC-sheets and EC-zippering. Thus, the present study, which was conducted in gel-excluded conditions, would provide an important starting point for understanding the mechanism of the heart and vein connection.

## ACKNOWLEDGEMENTS

We thank K. Kawakami (National Institute of Genetics) for the Tol2 system and N. Yamaguchi (University of Cambridge) for generating KI lines. We are grateful to M. Sone, H. Toyoshima, K. Hiratomi, and E. Hanimura for technical assistance, K. Shioya for fish care, and S. Fukuhara for helpful advice. This work was supported in part by the grants: JSPS KAKENHI (No. 19H01022 to N.M.; No. 21K15337 and No. 23K14224 to M.F.), AMED-CREST (No. 13414779 to N.M.), JST grant (JPMJPF2018 and JPMJPF2024-3 to N.M.) and Takeda Science Foundation to N.M.. This work was supported by COI-NEXT Support Unit for Imaging Science at Kento.

## AUTHOR CONTRIBUTIONS

Conceptualization, M.F. and N.M.; formal analysis, M.F.; investigation, M.F.; resources, M.F., H.F., and H.N.; Writing – Original Draft, M.F. and N.M.; Writing – Review & Editing, M.F., H.T., A.C., and N.M.; visualization, M.F.; supervision, N.M.; project administration, N.M.; funding acquisition M.F. and N.M.

## DECLARATION OF GENERATIVE AI AND AI-ASSISTED TECHNOLOGIES

We have not used AI

## INCLUSION AND DIVERSITY

We support inclusive, diverse, and equitable conduct of research.

## SUPPLEMENTAL INFORMATION TITLES AND LEGENDS

Figure S1. Timeline of Artery-Endo connection and Endo-sheet formation

Figure S2. Diversity of CCV-Endo adhesion mode

Figure S3. Endocardial ECs pull venous ECs along the anterior-posterior axis during EC-zippering

Figure S4. Gradual elongation of the endo-sheet

Figure S5. Gradual elongation of the entire embryo and the ventral tissue around the heart

Figure S6. Identification of Endo-sheet specific molecule

Figure S7. Cdh6 expression in various tissues

Figure S8. Cdh6-EGFP dynamics during initial contact and subsequent adhesion.

## SUPPLEMENTAL INFORMATION

Figures S1-8 and Video S1-20.

**Video S1. Time-lapse imaging of EC-zippering of a *Tg(fli1a:kikGR)* embryo from 60 hpf, related to Figure 1D**.

CCV-sheet (cyan) and Endo-sheet (orange) fuse in a zipper closing manner.

**Video S2. Time-lapse imaging of EC-zippering of a *Tg(kdrl:EGFP)* embryo from 60 hpf, related to Figure S2A.**

CCV-sheet and Endo-sheet fuse from the posterior edge toward the anterior edge.

**Video S3. Time-lapse imaging of EC-zippering of a *Tg(kdrl:EGFP)* embryo from 60 hpf, related to Figure S2A.**

CCV-sheet and Endo-sheet fuse from the anterior edge toward the posterior edge.

**Video S4. Time-lapse imaging of EC-zippering of a *Tg(kdrl:EGFP)* embryo from 60 hpf, related to Figure S2A.**

CCV-sheet and Endo-sheet fuse from both anterior and posterior edge toward the middle.

**Video S5. Time-lapse imaging of EC-zippering of a *Tg(kdrl:EGFP)* embryo from 60 hpf, related to Figure S2A.**

CCV-sheet and Endo-sheet fuse from both anterior and posterior edges toward the middle.

**Video S6. Time-lapse imaging of EC-zippering of a *Tg(kdrl:EGFP)* embryo from 72 hpf, related to Figure S2B.**

CCV-sheet and Endo-sheet fuse from both anterior and posterior edges toward the middle.

**Video S7. Time-lapse imaging of CCV-sheet and Endo-sheet connection of a *Tg(kdrl:EGFP)* embryo from 48 hpf, related to Figure S2B.**

Individual ccvEC randomly adheres to endoECs.

**Video S8. Time-lapse imaging of EC shape during EC-zippering of a *Tg(fli1a:LIFEACT-mCherry)* embryo from 60 hpf, related to Figure 2A**.

Cell contours during EC-zippering are visualized by LIFEACT-mCherry.

**Video S9. Time-lapse imaging of EC-zippering of a DMSO treated *Tg(fli1a:kikGR)* embryo from 60 hpf, related to Figure 4C**.

EC-zippering in normal condition.

**Video S10. Time-lapse imaging of EC-zippering of a Veramapamil-treated *Tg(fli1a:kikGR)* embryo from 60 hpf, related to Figure 4C**.

EC-zippers facilitated in the absence of heartbeat.

**Video S11. Time-lapse imaging of EC-zippering of a Isoprenaline treated *Tg(fli1a:kikGR)* embryo from 60 hpf, related to Figure 4C**.

EC-zippering inhibited in conditions of increased heartbeat.

**Video S12. Time-lapse imaging of Cdh6-EGFP at the zippering front of a *TgKI(cdh6-EGFP);Tg(fli1a:LIFEACT-mCherry)* embryo from 60 hpf, related to Figure 5C**.

Arrow points to the Cdh6-EGFP signal that could be detected at the adhesion front of EC-zippering.

**Video S13. Time-lapse imaging of Cdh6-EGFP in Endo-sheet of a *TgKI(cdh6-EGFP);Tg(fli1a:LIFEACT-mCherry)* embryo from 60 hpf, related to Figure 5D**.

Arrowheads point to the Cdh6-EGFP positive actin protrusions in Endo-sheet.

**Video S14. Time-lapse imaging of Cdh6-EGFP in CCV-sheet of a *TgKI(cdh6-EGFP);Tg(fli1a:LIFEACT-mCherry)* embryo from 60 hpf, related to Figure 5E**.

Arrows point to the stable linearized Cdh6-EGFP signal in CCV sheet.

**Video S15. Time-lapse imaging of a *Tg(kdrl:EGFP) cdh6^+/+^* embryo from 60 hpf, related to Figure 6C**.

Normal EC-zippering in *cdh6* expressing embryo.

**Video S16. Time-lapse imaging of a *Tg(kdrl:EGFP) cdh6^-/-^* embryo from 60 hpf, related to Figure 6C**.

A single EC elongates and closes the zipper.

**Video S17. Time-lapse imaging of a *Tg(kdrl:EGFP) cdh6^-/-^* embryo from 60 hpf, related to Figure 6C**.

A single ccvEC adhere to an endoEC but fails to close the zipper.

**Video S18. Time-lapse imaging of an isoprenaline-treated *Tg(kdrl:EGFP) cdh6^+/+^* embryo from 60 hpf, related to Figure 6G**.

Endo-sheet structure is maintained under conditions of increased heatbeat.

**Video S19. Time-lapse imaging of an isoprenaline-treated *Tg(kdrl:EGFP) cdh6^-/-^* embryo from 60 hpf, related to Figure 6G**.

EGFP-negative holes are detected in cdh6^-/-^ Endo-sheet under conditions of increased heatbeat.

**Video S20. Time-lapse imaging of a *TgKI(cdh5-EGFP);Tg(fli1a:LIFEACT-mCherry)* embryo from 60 hpf, related to Figure 7B**.

White arrowhead points to the low EGFP intensity region, and yellow arrowhead points to the high EGFP intensity region.

## STAR METHODS

### RESOURCE AVAILABILITY

#### Lead contact

Further information and requests for resources and reagents should be directed to and will be fulfilled by the lead contact, Naoki Mochizuki.

#### Materials availability

Further information and requests for resources and reagents listed in the key resources table should be directed to the lead contact.

### EXPERIMENTAL MODEL AND SUBJECT DETAILS

#### Animal studies

Zebrafish (*Danio rerio*) were maintained and bred in 28 °C water (pH 7.25 and conductivity 500 μS) with a 14 h on/10 h off light cycle. Embryos and larvae were incubated in the E3 medium at 28 °C. For confocal imaging, zebrafish embryos and larvae were dechorionated and anesthetized in 0.016% tricaine (Sigma-Aldrich, A5040) in the E3 medium. Animal experiments were approved by the institutional animal committee of the National Cerebral and Cardiovascular Center and performed according to the guidelines of the institute (Permit number: 2405) that follow the national (Japan) ethical and animal welfare regulations.

The fish lines used in this study were: AB as a wild-type line, *Tg(mly7:EGFP-CAAX)^ncv^*^54^*, Tg(fli1:myr-mCherry)^ncv^*^1^*, Tg(fli1:LIFEACT-mCherry)^ncv^*^7^*, TgBAC(dab2:EGFP)^ncv^*^67^*, Tg(fli1:kikGR) ^ncv^*^522^*, Tg(kdrl:EGFP) ^s^*^843^*, Tg(hsp70:zGrad)^ncv^*^561^*, Tg(fli1:LIFEACT-kikGR)^ncv^*^563^*, Tg(fli1:LIFEACT-HaloTag)^ncv^*^564^*, Tg(fli1:myl9b-HaloTag)^ncv^*^565^*, Tg(fli1:Golgi-HaloTag)^ncv5^*^66^*, TgKI(cdh6-EGFP)^ncv3^*^01^*, TgKI(cdh6-mCherry)^ncv3^*^04^, *TgKI(cdh5-EGFP)^ncv3^*^08^. Both males and females were used in this study.

### METHOD DETAILS

#### Plasmids

The Tol2 vector system was kindly provided by K. Kawakami (National Institute of Genetics, Japan).^39,40^

To construct the Tol2-hsp70-zGrad-pA plasmid, promoter/enhancer sequence of the hsp70 gene was obtained by PCR. zGrad fragment derived from pCS2+zGrad-pA (Addgene plasmid #119716; http://n2t.net/addgene: 119716 ; RRID:Addgene_119716) by PCR was inserted into the pTol2-hsp70 vector. Primers to amplify the *zGrad* were as follows: 5’-

ATCCACCGGTCGCCAatggagacggagatg-3’; 5’-

CTGCAGGAATTCGATttagctggagacggt-3’

To construct the Tol2-fli1a-lifeact-HaloTag-pA plasmid, promoter/enhancer sequence of the fli1a gene was obtained by PCR. HaloTag fragment derived from pC-Halo-N1 by PCR was inserted into the pTol2-fli1a-lifeact vector. Primers to amplify the *HaloTag* were as follows: 5′ - gaagaacgggatccaccggtcgccaccgaaatcggtactggc-3′ ; 5′ - atagttctagctcgagttaaccggaaatctccagagtagacagc-3′

To construct the Tol2-fli1a-myl9b-HaloTag-pA plasmid, promoter/enhancer sequence of the fli1a gene was obtained by PCR. HaloTag fragment derived from pC-Halo-N1 by PCR was inserted into the pTol2-fli1a-myl9b vector.

Primers to amplify the *HaloTag* were as follows: 5′ - ctggctttccattcgacccccattatgtgg-3′ ; 5′ - ccagtgaattatttaaccggaaatctccag-3′

To construct the pCS2+-EGFP-pA-cryaa-mCherry-pA plasmid, a promoter/enhancer sequence of the *cryaa* gene obtained by PCR, an EGFP-pA fragment derived from pEGFP-N1 (Clontech), a mCherry-pA fragment derived from pmCherry-N1 (Clontech) were inserted into the pCS2+ccvECtor. Primers to amplify the *cryaa* promoter were as follows: 5′ - gagctgtacaagtaaaggtaccatcgatga-3′ ; 5′ - TTATGATCTAGAGTCCTACTTGTACAGCTC-3′.

To construct the pCS2+cdh6-EGFP-pA-cryaa-mCherry-pA plasmid and pCS2+cdh6-mCherry-pA-cryaa-EGFP-pA, a genomic fragment spanning the 329 bp upstream of *cdh6* exon 11 through the end of exon11 obtained by PCR was inserted into the pCS2+-EGFP-pA-cryaa-mCherry-pA plasmid or pCS2+-mCherry-pA-cryaa-EGFP-pA plasmid (exon numbering is based on transcript ID: ENSDART00000131506.3). Primers to amplify the cdh6 genome fragment were as follows: 5′ - ggtcgccaccaatgggtggatggac -3′ ; 5′ - ccttgctcacggagtcccaatcgct -3′

To construct the pCS2+cdh5-EGFP-pA-cryaa-mCherry-pA plasmid, a genomic fragment spanning the 491 bp upstream of *cdh5* exon 11 through the end of exon11 and exon 12 genomes obtained by PCR was inserted into the pCS2+-EGFP-pA-cryaa-mCherry-pA plasmid (exon numbering is based on transcript ID: ENSDART00000108766.3). Primers to amplify the cdh5 genome fragment were as follows: 5′ -ctgtatcattaccatcctggtaatcgtaatcctca-3′ ; 5′ - ccatgccagcgtgtgttaatttaactggaattc-3′ ; 5′ - ctgtatcattaccatcctggtaatcgtaatcctca -3′ ; 5′ - ccatggactgatatcgtaggagctatccga-3′

#### Transgenic zebrafish lines

We used the following previously published zebrafish lines: *Tg(mly7:EGFP-CAAX)^ncv^*^54^*, Tg(fli1:myr-mCherry)^ncv1^, Tg(fli1:LIFEACT-mCherry)^ncv7^, Tg(fli1:kikGR) ^ncv5^*^22^*, Tg(kdrl:EGFP) ^s8^*^43^. The *Tg(hsp70:zGrad)^ncv5^*^61^*, Tg(fli1:LIFEACT-kikGR)^ncv563^, Tg(fli1:LIFEACT-HaloTag)^ncv564^, Tg(fli1:myl9b-HaloTag)^ncv565^, Tg(fli1:Golgi-HaloTag)^ncv566^* lines were established by injecting transposase mRNA (25 pg) and either Tol2-based plasmids (30 pg) into one-cell stage AB. Tol2 transposase mRNAs were *in vitro* transcribed with SP6 RNA polymerase from the linearized pCS2-TP vector using the mMESSAGE mMACHINE kit (Thermo Fisher Scientific, AM1340). Throughout the text, all Tg lines used in this study are simply described without their line numbers. For example, *Tg(myl7:GFP-CAAX) ^ncv54^* is abbreviated to *Tg(myl7: GFP-CAAX)*. In addition, *cryaa*:mCherry is omitted throughout the main text: *TgKI(cdh6-EGFP, cryaa:mCherry)^ncv301^* is written as *TgKI(cdh6-EGFP)*.

#### Generation of knockout zebrafish by CRISPR/Cas9 system

*cdh6^ncv147^* knockout line was established by the following method. gRNAs targeting *cdh6 5’UTR and 3’UTR* were designed using CRISPR direct software.^41^ Embryos, co-injected with 0.5 nl of a KO cocktail consisting of gRNA (final concentration 12µM) and Cas9 protein tagged with a nuclear localization sequence (IDT) (final concentration 2 ng/nl) at the on e-cell stage, were raised to adulthood and crossed with wild-type AB to identify germline-mutated founders.

Screening for founders was conducted by genomic PCR and subsequent sequencing using the following primer sets: 5′ - ggctattgtgaccgcgcatgctga-3′ and 5′ -gcttacaaattggcaaacca-3′. For the genotyping of the mutants, PCR analyses of genomic DNAs were routinely performed using the following primer sets: 5′ -ggctattgtgaccgcgcatgctga-3′ and 5′ -atgcttcccttcctaattaaaggc-3′ and 5′ - ggctattgtgaccgcgcatgctga-3′ and 5′ -gcttacaaattggcaaacca-3′.

#### Generation of knock-in zebrafish by CRISPR/Cas9 system

*TgKI(cdh6-EGFP)^ncv3^*^01^, *TgKI(cdh6-mCherry)^ncv3^*^04^, and *TgKI(cdh5-EGFP)^ncv308^*knock-in lines were generated following the published method.^42^ A gRNA target site was designed to target both the donor plasmid, thereby linearizing it in the intron, and the endogenous genomic intron.

Early one-cell-stage embryos were injected with 0.5 nl of a KI cocktail consisting of gRNA (final concentration 12 µM), donor plasmid (final concentration 6 pg/nl) Cas9 protein tagged with a nuclear localization sequence (IDT) (final concentration 2 ng/nl), and phenol red (final concentration 0.05%). Embryos were visually screened at 3 dpf for fluorescence as described below.

#### Genotyping

Genome DNA was extracted from adult fish fins or whole embryos. Tissues were dipped into 300 µl NaOH(1 M, FUJIFILM, 198-13765) and boiled for 5min. 3 µl Tris-HCl (0.5 M pH8.0, SIGMA, T1503) was added and centrifuged at 13500 rpm for 10 min. 1 µl of the supernatant was used as template DNA for PCR. The specific genome regions were amplified by KOD Fx Neo (TOYOBO, KFX-201) using the following primer sets.

For wild type allele 5′ -ggctattgtgaccgcgcatgctga-3′ and 5′ - atgcttcccttcctaattaaaggc-3′

For *cdh6* KO allele 5′ -ggctattgtgaccgcgcatgctga-3′ and 5′ - gttctgtaatttattttcagtgta-3′

#### Chemical treatment

Fish were treated with the following chemicals: Isoprenaline hydrochloride (300 µM, Signa-Aldrich, I5627) dissolved in fish water containing dimethyl sulfoxide (DMSO, 0.2%, FUJIFILM, 045-24511) immediately before treatment, to promote heatbeat; Verapamil hydrochloride (200 μM, Wako, 222-00781) dissolved in fish water containing Methanol (0.1%, FUJIFILM, 137-01823) immediately before treatment to repress heatbeat.

#### Heat shock

*TgKI(cdh6-EGFP);Tg(hsp70:zGrad);(fli1a:LIFEACT-mCherry)* and *TgKI(cdh5-EGFP);Tg(hsp70:zGrad);(fli1a:LIFEACT-mCherry)* embryos were heat-shocked at 58 hpf for 30 min at 39.5 °C. Images were obtained before and 6 hours after heatshock (Figure 6H-L, 7D-G). After imaging, each embryo was genotyped by PCR using the following primer sets.

For zGrad 5′ - atccaccggtcgccaatggagacggagatg-3′ and 5′ - ctgcaggaattcgatttagctggagacggt-3′

For *cdh6* wild type allele 5′ -gggtgttgggtttatttatgga -3′ and 5′ - aggaagatgtcaccgcagat-3′

For *cdh6* knock-in allele 5′ - gggtgttgggtttatttatgga-3′ and 5′ - gctgaacttgtggccgtttacgtc-3′

For *cdh5* wild type allele 5′ - ccccagagcattttgaagtc -3′ and 5′ - gatgaacctacccaggatgg -3′

For *cdh5* knock-in allele 5′ - cagcaactgttttgacaccaa -3′ and 5′ - gctgaacttgtggccgtttacgtc -3′

#### Imaging

To obtain images of embryos without pigmentation, a 0.2 mM solution of N-phenylthiourea (PTU) (Tokyo Chemical Industry, P0237) was added to the E3 medium from 24 hpf. Embryos and larvae were anaesthetized with 0.1mg/ml of tricaine (Sigma-Aldrich, A5040) and then mounted in 1% low-melting agarose (Sigma-Aldrich, A9414). For time-lapse imaging, agarose gel around the ventral tissue was excluded (Figure S4B, right).

Confocal imaging was performed using an FV1200 confocal microscope (Olympus), and an FV3000 confocal microscope (Olympus). We used a 20x water immersion objective (Olympus XLUMPlan FL_N, 1.0 N.A.). All confocal images were processed and analyzed with Imaris 9.8.0, 9.9.1, or 10.0.0 software (Oxford Instruments).

#### Imaging of HaloTag-labeled protein

To obtain images of *Tg(fli1:LIFEACT-HaloTag)^ncv564^, Tg(fli1:myl9b-HaloTag)^ncv565^, Tg(fli1:Golgi-HaloTag)^ncv566^*, embryos carrying cristallin:TagBFP at the 48 hpf were placed into 1.5 ml tubes, and most of the E3 buffer was removed such that embryos were just covered sufficiently. 100 µl of JFX554 (10nM, Promega, HT1030) or JFX650 (10nM, Promega, HT107A) dye solution was then added to the tubes for more than 30 min. After pre-staining, zebrafish were directly embedded into 1% agarose gel for imaging. For timelapse imaging, 8 ml of 10 nM HaloTag-JFX554 or HaloTag-JFX650 dye containing E3 buffer was added to the glass bottom dish.

#### Photoconversion, photoactivation and laser ablation

Photoconversion was performed using 405 nm laser line around 2-3% laser power. Experiments were conducted using single-photon hybrid detectors for visual control with 488 nm excitation of KikmeGreen and 561 nm excitation for KikmeRed. Photoactivation of PA-GFP was performed using 405 nm laser line around 2-3% laser power. Laser ablation was performed using 920 nm laser line at 20-30% laser power. Experiments were conducted using two-photon hybrid detectors for visual control with 920 nm excitation of kdrl:EGFP.

#### Reanalysis of single-cell RNA sequence

Processed scRNA-seq (x10) data for 24 hpf zebrafish embryos from a previous study were obtained from the web-based resource (www.tinyurl.com/scZfish2018). Expression profiles of cells were clustered using the Single-Cell Analysis in Python (ScanPy) version 1.7.0. We removed genes that were not expressed in at least 3 cells and cells that did not have at least 200 detected genes. We removed cells with more than 6,000 unique molecular identifiers (UMIs) or more than 20,000 leads. The data was then normalized using size factor normalization, such that every cell has 10,000 counts and log transformed. We computed highly variable genes using default parameters and then scaled the data to a maximum value of 10. We then extracted *etv2*-expressing cells (log-transformed expression levels > 0.5) and computed principal component analysis. Cells were clustered by constructing a shared nearest neighbor graph and clusters were visualized by umap projection. We annotated each cluster using *fn1a* and *gata5* for endotcardial endothelial lineage (Figure S5). Endocardial endothelial cells were then subclustered, constructing a shared nearest neighbor graph and clusters were visualized by umap projection. We annotated endocardial sheet endothelial cells using *dab2* since *dab2* promoter was specificaly activite in endocardial sheet using *TgBAC(dab2:dEGFP)* line. Endocardial sheet endothelial cell-specific adhesion molecules were explored by a comprehensive comparison of the clusters. Clusters of other tissues were annotated using the default tissue ID (Figure S6).

#### Quantitative PCR (qPCR)

Total RNA was prepared from 2.5 dpf embryos using the TRIzol Reagent (Thermo Fisher Scientific 15596026) according to the manufacturer’s instructions. RNAs were reverse-transcribed with oligo dT primers using PrimeScript FAST RT regent kit with gDNA Eraser (TAKARA, RR092S). The PCR reaction was performed using TB Green Premix Ex Taq2 FAST qPCR (TAKARA, RR830S) in a CFX Connect thermal cycler (Bio-Rad). The primers used for qPCR are listed below. The qPCR results were normalized to *beta-actin* expression. *cdh6*, 5′ -AAATGTGCAATGCTGAGGCG-3′ and 5′ - GCACCACGATCAAAAGCAGG-3′ ; *beta-actin*, 5′ - GATCTTCACTCCCCTTGTTCA-3′ and 5′ -GGCAGCGATTTCCTCATC-3′.

### QUANTIFICATION AND STATISTICAL ANALYSIS

#### Counts of time of first connection

For the quantification of Figure 1C, the *Tg(kdrl;EGFP)* embryos were observed under SZX16 stereo microscope (Olympus) every 12 hours from 36 hpf to 108 hpf. When lateral CCVs are adhered to the Endo-sheet, the embryo was counted as a connected embryo. The percentage of the connected embryo out of the entire embryo number was calculated at each time point and plotted as histogram.

#### Zipper length measurement

For quantification of Figures 1E, 4D, and 6E, images of *Tg(kdrl;EGFP)* embryos were recorded using an FV3000 confocal microscope (Olympus). EGFP+ length from the posterior edge to the zipper front was measured using Fiji ImageJ software every 1 hour for Figures 1E and 4D, and at 10 h for Figure 6E.

#### Counts of adhesion cell number

For quantification of Figure 1F, *Tg(kdrl;EGFP)* embryos were recorded using an FV3000 confocal microscope (Olympus). The EC nuclear number labelled by strong EGFP signaling in the front layer of the adhesion site was counted at the end point of zippering.

#### Cell boundary analysis

For analysis of Figures 2B and 2E, images of *Tg(fli1a:LIFEACT-mCherry)* embryos were obtained by FV3000 confocal microscope (Olympus). Cell counter was manually extracted as mCherry-positive signal every 30 min and projected manually.

#### Kymograph

Cell boundary dynamics at the initial adhesion for Figure 2C were analyzed using Videos of the *Tg(fli1a:LIFEACT-mCherry)* line. Positional changes of the endoEC and ccvEC cell boundary were quantified and grouped in 10 to 30-minute bins. T=0 indicates the first frame in which endoEC and ccvEC made contact. Kymographs were generated using the ‘KymoRescliceWide’ FIJI ImageJ plugin with a linewidth of 1 pixel. Cell boundary was manually traced from the kymographs. The track’s y coordinates were converted into time (h) and plotted.

#### Cell polarity analysis

Cell polarity dynamics during zippering for Figure 2F were analyzed using snapshots of the 50% zipper closed *Tg(kdrl:EGFP);(fli1a:Golgi-HaloTag)* line. The position of Golgi and nucleus were automatically extracted as x and y coordinates using the ‘spot’ function of Imaris software (Oxford Instruments). Cell polarities were quantified as Golgi-to-nucleus axis in each ECs. The frequency was represented as a polar histogram every 30 degrees.

#### KikmeRed+ region aspect ratio measurement

For quantification of Figures 3B, 3H, and 4B, images of photoconverted *Tg(fli1a:kikGR)* embryos were obtained by FV3000 confocal microscope (Olympus). The KikmeRed+ region was extracted using Fiji ImageJ software and the aspect ratio was measured by shape descriptors analysis every 12 hours for Figure 3B, and at 2 dpf and 3 dpf for Figures 3F and 4B. *Tg(fli1a:LIFEACT-kikGR)* was also used for Figure 3B.

#### Cell shape measurement

For quantification of Figure 3C, images of photoconverted *Tg(fli1a:kikGR)* embryos were obtained by FV3000 confocal microscope (Olympus) at 2 dpf and 3 dpf. Two separated EC in the Endo-sheet were photoconverted at 2 dpf. The KikmeRed+ region was extracted using Fiji ImageJ software and the aspect ratio was measured by shape descriptors analysis.

#### Cell size measurement

For quantification of Figure 3D, images of photoconverted *Tg(fli1a:kikGR);(fli1a:LIFEACT-HaloTag)* embryos were obtained by FV3000 confocal microscope (Olympus) at 2 dpf and 3 dpf. The entire Endo-sheet was photoconverted at 2 dpf. Cell counter was manually extracted as HaloTag-positive signal. Cell size was measured by shape descriptors analysis using Fiji ImageJ software.

#### Myosin accumulation measurement

For quantification of Figure 3E, images of photoconverted *Tg(fli1a:kikGR);(fli1a:myl9b-HaloTag)* embryos were obtained by FV3000 confocal microscope (Olympus) at 2 dpf and 3 dpf. The entire Endo-sheet was photoconverted at 2 dpf. Myl9b accumulation was measured by signal intensity measurement of 10 randomly chosen ROI using Fiji ImageJ software.

#### Convergent extension measurement

For quantification of Figure 3F and 3G, 100 ng of PA-GFP mRNA was injected at 1 cell stage. Five points (one was set at the center, and the other four were 100 µm from the center point.) of the ventral tissue of 48 hpf embryos were labeled by photoactivation with FV3000 confocal microscope (Olympus). The images of PA-GFP labelled diamond were obtained every 24 hours from 2 dpf to 4 dpf. The xy coordinates of the center of each PA-GFP points were gained using Fiji ImageJ software and the aspect ratio of the PA-GFP labelled diamond was measured.

#### Cell polarity projection

For the qualification of Figure 3J and 3K, images of *Tg(fli1a:kikGR);(fli1a:Golgi-HaloTag)* embryos were recorded using an EV3000 confocal microscope (Olympus) at 2 dpf and 3 dpf. EndoECs were photoconverted to kikumeRed at 2 dpf. Golgi-to-nucleus axis was measured using the ‘spot’ function in Imaris software (Oxford Instruments). Nuclei whose Golgi-to-nucleus axis is 45-135 degrees were labeled as blue, 225-315 degrees as pink, and 135-225 and 315-45 as yellow. Cell polarity mapping of individual embryos was merged using Fiji ImageJ software.

#### Zipper speed measurement

For quantification of Figure 4E, images of *Tg(kdrl;EGFP)* embryos were recorded using an FV3000 confocal microscope (Olympus). EGFP+ length from the posterior edge to the zipper front was measured using Fiji ImageJ software every 1 hour. The zippering speed for every 2 hours was calculated from 0 h to 10 h and averaged in each sample.

#### Cdh6-EGFP signal intensity measurement

For quantification of Figure 5F, images of *TgKI(cdh6-EGFP);Tg(fli1a:LIFEACT-mCherry)* embryos were recorded using an FV3000 confocal microscope (Olympus). Images were filtered by a Median filter. EGFP signal at the cell boundary was measured using Fiji ImageJ ‘line’ function before and 6 hours after Verapamil treatment.

#### EC moving distance measurement

For quantification of Figure 6F, *Tg(kdrl;EGFP)* embryos were recorded using an FV3000 confocal microscope (Olympus). EC nuclei labelled by strong EGFP signal were tracked using Imaris software (Oxford Instruments). The x and y coordinates were calculated every 30 minutes by Imaris software (Oxford Instruments) as displacement length x and y respectively.

#### Count of hole number in Endo-sheet

For quantification of Figures 6G and 6H, images of *Tg(kdrl;EGFP)* embryos were recorded using an FV3000 confocal microscope (Olympus). Images at 1 hour after isoprenaline treatment were converted to binary image using Fiji ImageJ. The white signal completely surrounded by the black signal was counted as a hole.

#### endoEC size measurement

For quantification of Figures 6K and 7E, images of *TgKI(cdh6-EGFP);Tg(hsp70:zGrad);(fli1a:LIFEACT-mCherry)* or *TgKI(cdh5-EGFP);Tg(hsp70:zGrad) ;(fli1a:LIFEACT-mCherry)* embryos were obtained by FV3000 confocal microscope (Olympus) at 0h and 6h post heat shock. Each cell contour was visualized by LIFEACT-mCherry signals. 10 cells were randomly picked up per individual and cell sizes were measured using Fiji ImageJ software.

#### Endo-sheet A-P length measurement

For quantification of Figures 6J and 7D, images of *TgKI(cdh6-EGFP);Tg(hsp70:zGrad);(fli1a:LIFEACT-mCherry)* or *TgKI(cdh5-EGFP);Tg(hsp70:zGrad) ;(fli1a:LIFEACT-mCherry)* embryos were obtained by FV3000 confocal microscope (Olympus) at 0h and 6h post heat shock. The Endo-sheet length from the posterior edge to the anterior edge labelled by LIFEACT-mCherry was measured by using Fiji ImageJ software.

#### Data analysis and statistics

Data were analyzed using Excel (Microsoft), Matplotlib, and Plotnine and were presented as individual data or mean ± s.d. Sample numbers were indicated in the figure legends. Statistical analysis was performed by paired samples *t*-test (Figures 3A, 3C, 3D, 3E, 6L, 7E, 7F), Welch’s *t*-test (Figures 3G, 3I, 4B, 4D, 4E, 5G), two-tailed Student’s *t*-test (Figures 6E). Statistical significance was defined as p < 0.05.

