## Supplemental information for "Endothelial-zippering proceeds by sensing heartbeat-driven force through Cadherin-6 during heart-vessel connection in zebrafish"

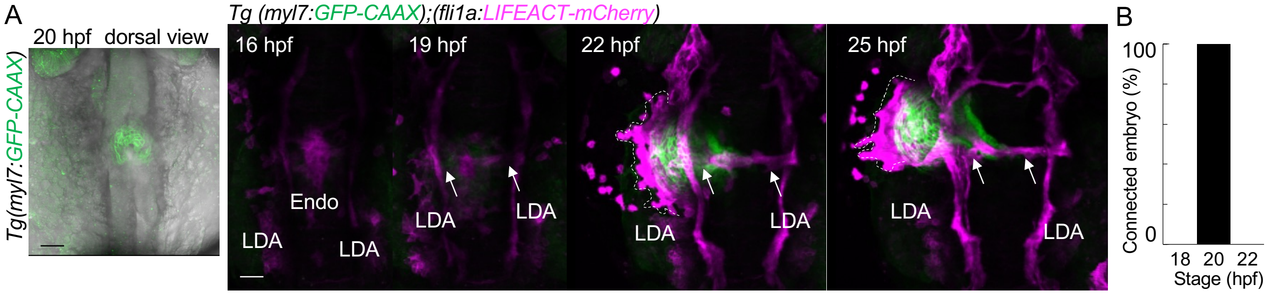


**Figure S1. Timeline of Artery-Endo connection and Endo-sheet formation**

(A) Representative confocal images of the dorsal view of *Tg(myl7:GFP-CAAX)* embryo at 20 hpf (left). Representative time-sequential images of *Tg(myl7:GFP-CAAX);(fli1a:LIFEACT-mCherry)* embryo from 16 hpf. The dashed line outlines the Endo-sheet. Arrows point to the LDA-Endo connection established during 19-22 hpf.

(B) Graph shows the percentage of LDA-Endo connected embryos. *n*=5.

Scale bars 30 µm; LDA, lateral dorsal aorta; Endo, endocardium.

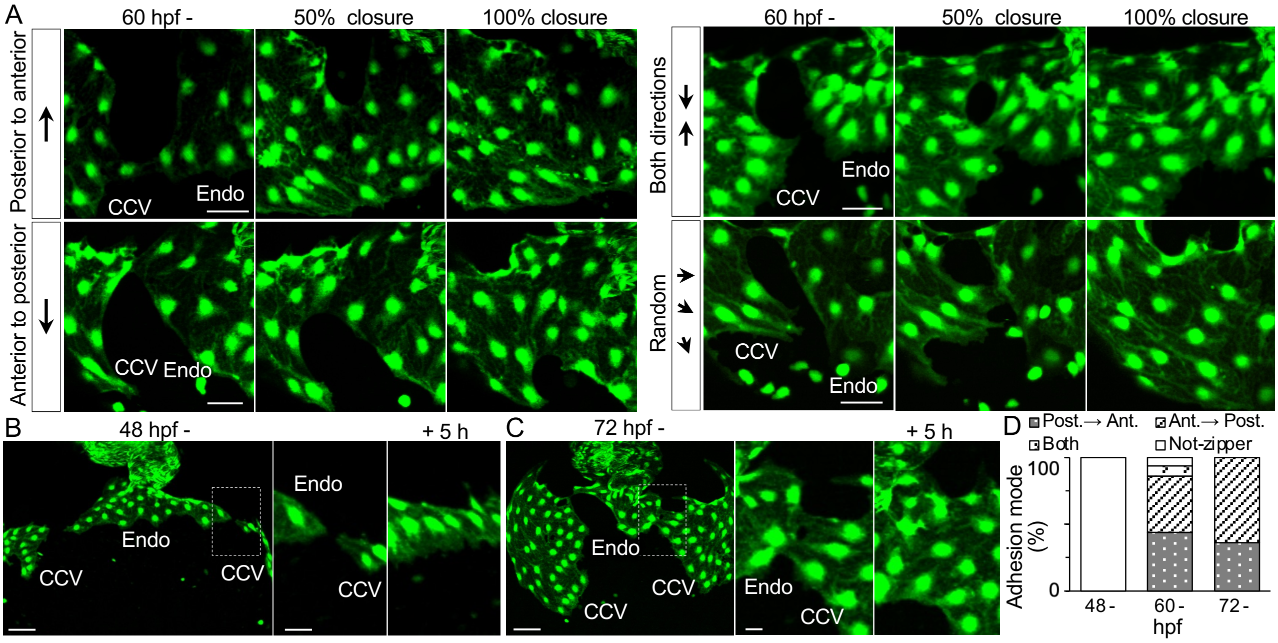


**Figure S2. Diversity of CCV-Endo adhesion mode**

(A) Representative time-sequential images of the ventral view of *Tg(kdrl:EGFP)* embryo from 60 hpf of four different adhesion directions from the initial adhesion site.

(B) Representative confocal images of the ventral view of *Tg(kdrl:EGFP)* at 48 hpf (left). The dashed box region at 48 hpf and 53 hpf are enlarged on the right.

(C) Representative confocal images of the ventral view of *Tg(kdrl:EGFP)* at 72 hpf (left). The dashed box region at 72 hpf and 77 hpf are enlarged on the right.

(D) Graph shows the percentage of four adhesion modes at three different stages. 48 hpf-;*n*=14, 60 hpf-;*n*=50, 72 hpf-;*n*=11.

Scale bars, 30 µm (A), 50 µm (B and C left), 20µm (B and C right); Endo, endocardium; CCV, common cardinal vein.

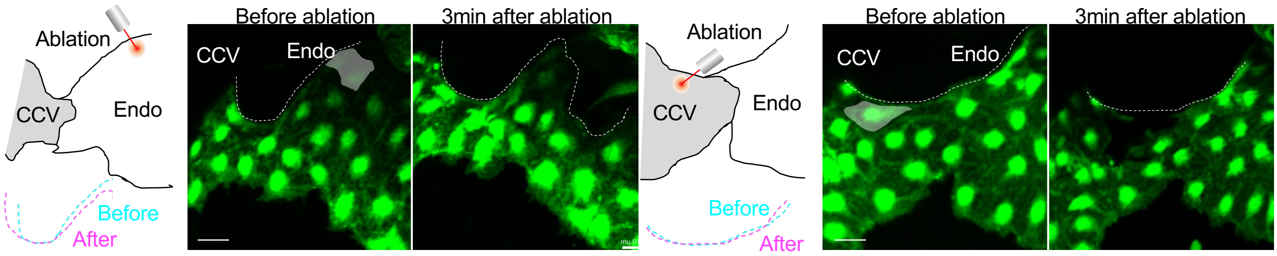


**Figure S3. Endocardial ECs pull venous ECs along the anterior-posterior axis during EC-zippering**
Schematic images of laser ablation (each panel left). The anterior endoEC (masked white) was laser ablated at 50 % zipper closure phase (left-panel centre). ). The posterior ccvEC (masked white) was laser ablated at 50 % zipper closure phase (right-panel centre). The white dashed lines outline the zipper front in each image. The axis of the zipper front was merged as cyan- (before ablation) and magenta- (after ablation) dashed lines. *n*=3 each.
Scale bars, 20 µm (A); Endo, endocardium; CCV, common cardinal vein.

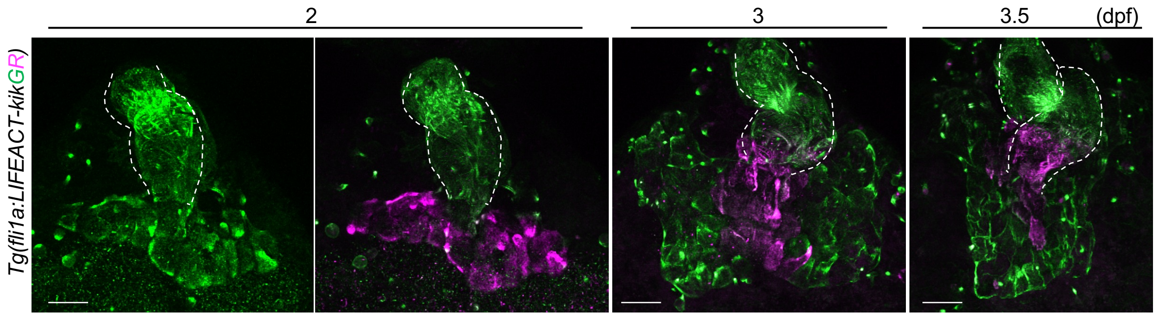


**Figure S4. Gradual elongation of the endo-sheet**

Representative snap-shot images of *Tg(fli1a:LIFEACT-KikumeGR)* embryo from 2 dpf to 3.5 dpf. The dashed line outlines the heart. Endo-sheet photoconverted into magenta. Scale bars; 10μm.

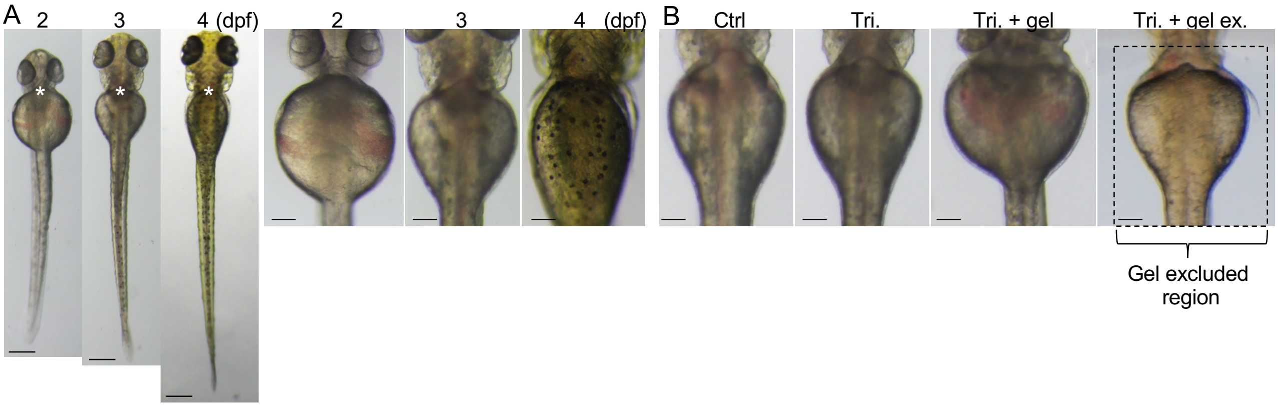


**Figure S5. Gradual elongation of the entire embryo and the ventral tissue around the heart**
(A) Ventral view of the entire embryo (left). * represent the heart-developing region enlarged on the right.
(B) Ventral view of the ventral tissue at 3 dpf in various culture conditions.
Scale bars, 200 μm (A left), 100 μm (A right and B); Tri, tricaine; gel, gel embedded; gel ex. gel excluded.

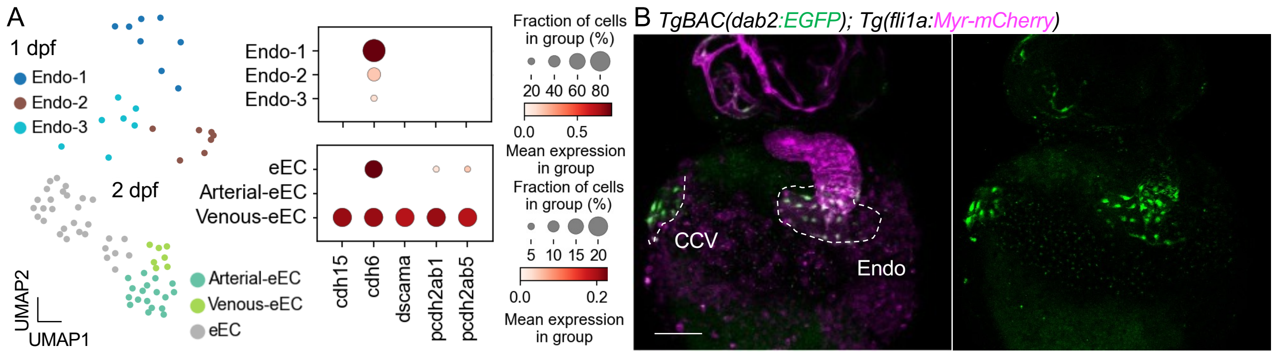


**Figure S6. Identification of Endo-sheet specific molecule**

(A) scRNA-seq. analysis of 1 dpf and 2 dpf embryos. eEC data annotated by high fn1a, gata5 expression were subclustered. Dot plot shows the expression levels of Endo-sheet specific adhesion candidates.

(B) Representative confocal images of the ventral view of *TgBAC(dab2:EGFP);Tg(fli1a:Myr-mCherry)* embryo at 36 hpf.

Scale bar 100 μm; Endo, endocardium; CCV, common cardinal vein.

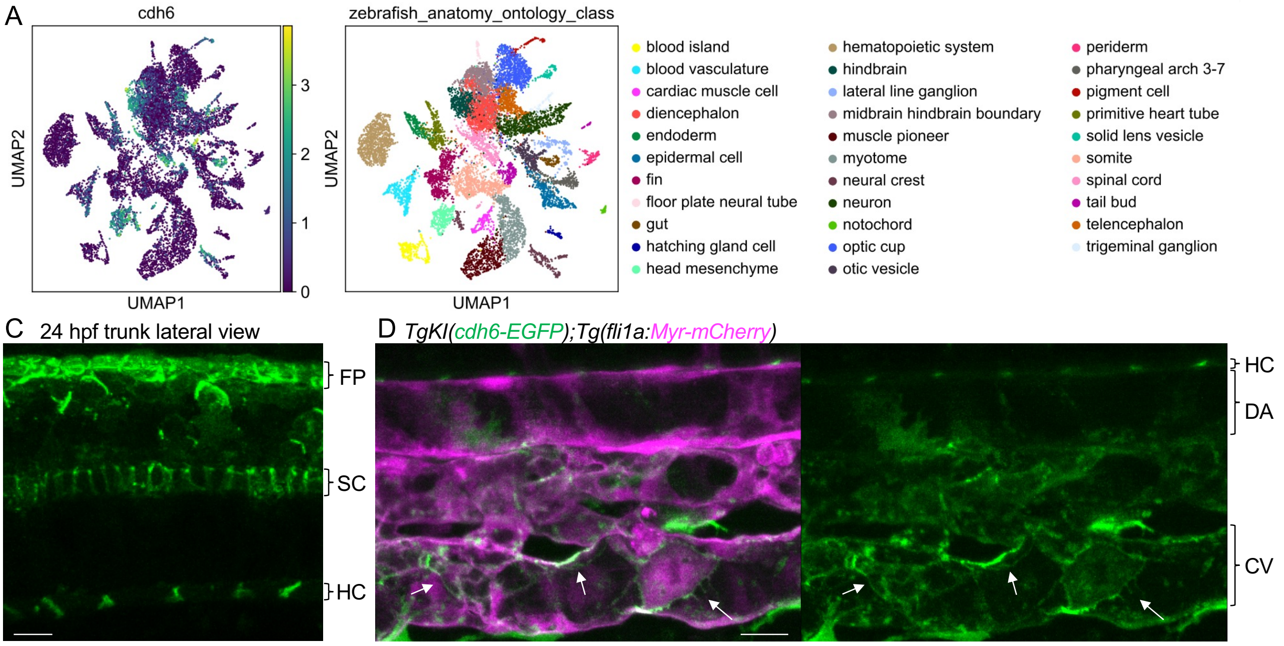


**Figure S7. Cdh6 expression in various tissues**

(A) scRNA-seq. analysis of 24 hpf embryo. UMAP-plot shows the *cdh6* mosaic expression pattern among multiple clusters (left). Clusters were annotated into each tissue (right).

(B) High magnification confocal images of the lateral view of the trunk of *TgKI(cdh6-EGFP)* trunk at 24 hpf.

(C) High magnification confocal images of the lateral view of *TgKI(cdh6-EGFP);Tg(fli1a:Myr-mCherry)* embryo at 48 hpf. Arrows point to the weak but specific Cdh6-EGFP signals in venous ECs.

Scale bars 50 μm (C),15 μm (D); FP, floorplate; SC, spinal cord; HC, hypochord; DA, dorsal aorta; CV, caudal vein.

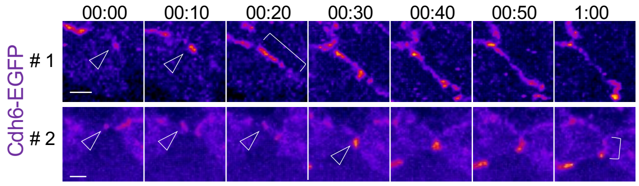


**Figure S8 Cdh6-EGFP dynamics during initial contact and subsequent adhesion.**
Ten-minute interval heatmap images of cdh6-EGFP of 2 embryos. Arrowheads point to the accumulation of Cdh6-EGFP. The intensity of EGFP is displayed as heatmap.
